# Age-related late-onset disease heritability patterns and implications for genome-wide association studies

**DOI:** 10.1101/349019

**Authors:** Roman Teo Oliynyk

## Abstract

**Background:** Genome-wide association studies and other computational biology techniques are gradually discovering the causal gene variants that contribute to late-onset human diseases. After more than a decade of genome-wide association study efforts, these can account for only a fraction of the heritability implied by familial studies, the so-called “missing heritability” problem.

**Methods:** Computer simulations of polygenic late-onset diseases in an aging population have quantified the risk allele frequency decrease at older ages caused by individuals with higher polygenic risk scores becoming ill proportionately earlier. This effect is most prominent for diseases characterized by high cumulative incidence and high heritability, examples of which include Alzheimer’s disease, coronary artery disease, cerebral stroke, and type 2 diabetes.

**Results:** The incidence rate for late-onset diseases grows exponentially for decades after early onset ages, guaranteeing that the cohorts used for genome-wide association studies overrepresent older individuals with lower polygenic risk scores, whose disease cases are disproportionately due to environmental causes such as old age itself. This mechanism explains the decline in clinical predictive power with age and the lower discovery power of familial studies of heritability and genome-wide association studies. It also explains the relatively constant-with-age heritability found for late-onset diseases of lower prevalence, exemplified by cancers.

**Conclusions:** For late-onset polygenic diseases showing high cumulative incidence together with high initial heritability, rather than using relatively old age-matched cohorts, study cohorts combining the youngest possible cases with the oldest possible controls may significantly improve the discovery power of genome-wide association studies.

## Introduction

Throughout the ages, late-onset diseases (LODs) were considered the bane of the lucky few who survived to an advanced age. Over the last couple of centuries, however, continuous improvements in sanitation, life and work environments, vaccinations, disease prevention, and medical interventions have extended the average life expectancy by decades.

With a growing fraction of the population being of advanced age, the leading causes of mortality are now heart disease, cancer, respiratory disease, stroke, and notably Alzheimer’s disease and other dementias (Murphy et al., 2017). The need—and with it, the effort being made—to determine the causes of late-onset diseases is ever increasing, and one of the targets of medicine has become combating aging itself in addition to specific age-related diseases (Franceschi et al., 2018).

One of the major goals of computational biology is to identify gene variants that lead to increased odds of late-onset diseases. Nevertheless, polygenic LODs remain resistant to the discovery of sufficient causal gene variants that would allow for accurate predictions of an individual’s disease risk (Manolio et al., 2009; Clarke and Cooper, 2010; Kumar et al., 2016). This is despite the fact that LODs with varied symptoms and phenotypes show high heritability in twin and familial studies (Zaitlen and Kraft, 2012).

At a young age, the human organism usually functions as well as it ever will. With time, the organism’s functions decline, leading to the common image of aging as one of thinning hair and a loss of pigmentation in what remains, increased wrinkling and altered pigmentation of the skin, reductions in height, muscle and bone mass, joint pain, and deficits in hearing, sight and memory (Fedarko, 2018). The combination of genetic liability, environmental factors, and the physiological decline of multiple organism systems leads to individual disease presentation. Genetic variation may be either protective or detrimental when compared to the average distribution of common gene variants that defines human conditions as it applies to polygenic LODs.

Researchers engaged in genome-wide association studies (GWASs) often set an unrealistic expectation that a combination of causal single nucleotide polymorphisms (SNPs)—also known as a polygenic score—will, irrespective of the patient’s age, completely predict an individual’s predisposition to an LOD to a degree matching the maximum heritability found in familial studies (Naj and Schellenberg, 2017; Silva et al., 2015). The lost heritability debate, in the case of LODs, often treats polygenic LODs as if they were binary hereditary phenotypic features rather than facets of failure processes that arise in the human body (Oh et al., 2014) when it is past its reproductive prime and when evolutionary selection is significantly relaxed compared to younger ages (Fedarko, 2018).

GWAS can implicate a subset of SNPs that can typically explain between 10 and 20% of the genetic heritability of a polygenic LOD (Visscher et al., 2017). There are two complementary hypotheses explaining this so-called missing heritability (Eyre-Walker, 2010; Yang et al., 2012; Thornton et al., 2013; Agarwala et al., 2013). The first is the hypothesis that LODs are caused by a combination of a large number of relatively common alleles of small effect (Goldstein, 2009). GWASs have been able to discover only a small number of moderate-effect SNPs, but a large number of SNPs remain below GWASs’ statistical discovery power. The second hypothesis states that LODs are caused by a relatively small number of rare, moderate- or high-effect alleles with a frequency below 1% that likely segregate in various proportions into sub-populations or families (Dickson et al., 2010; North and Beaumont, 2015) and are also under the radar of GWASs’ discovery power.

Both scenarios can contribute to observational facts, but their relative weights vary depending on the genetic architecture of an LOD (Park et al., 2011). Rare highly detrimental alleles become indistinguishable in their presentation from the OMIM cataloged conditions and will likely be diagnosed as a separate disease or syndrome. The population age distribution and individual disease progression of polygenic LODs are best understood by considering the aging process itself as an ongoing loss of function, which can be modulated by the genetic liabilities resulting from both common and rare SNP distributions combined with changing environmental and lifestyle variables. It has been determined (Anderson et al., 2011; Yang et al., 2015) that common variants very likely explain the majority of heritability for most complex traits.

While the findings of GWASs can explain only a fraction of heritability, the systematically collected SNP correlations provide a good indication of what to expect regarding the effect sizes and allele frequency distribution of as yet undiscovered SNPs (Eyre-Walker, 2010). Many studies focus on constructing hypotheses, defining the types of gene variants that could explain the missing heritability, proving why these gene variants are difficult to discover, and identifying the evolutionary processes that led to the hypothesized and observed gene variant distributions (Manolio et al., 2009; Clarke and Cooper, 2010; So et al., 2011; Zaitlen and Kraft, 2012; Thornton et al., 2013; Wood et al., 2014). These studies explore the effect sizes and allele frequencies that GWAS would expect to find for LODs as well as the genetic architecture of complex traits and their implications for fitness.

The age-related heritability decline of some LODs has been assumed for decades. The precise magnitude of heritability change with age is typically unknown for most LODs, and the effects are not understood and often ignored or overlooked. Most GWASs recommend homogeneity in cohort age—that is, that the same age window should be targeted–—although it has been suggested (Li and Meyre, 2013) that individuals with an early age of onset are likely to have greater genetic susceptibility. Discussing a replication study design, Li and Meyre (2013) stated, “Once the risk of false positive association has been ruled out by initial replication studies, the focus of the association can be extended to different age windows.” Another common approach is to “age adjust” the effect (Zaitlen et al., 2012) with the goal of removing or averaging out the effect of aging rather than examining its consequences more thoroughly. Two recent studies (Lin et al., 2014; Bjørnland et al., 2018) emphasize the need to explore “extreme phenotype sampling” in order to improve GWAS discovery, including using cohorts that are diverse in age.

One of the first geneticists to build a conceptual foundation for disease susceptibility, and the pioneer of the liability threshold approach, was D. S. Falconer in his studies of inheritance estimated from the prevalence among relatives (Falconer, 1965) and his 1967 follow-up study exploring the prevalence patterns of LODs, specifically diabetes (Falconer, 1967), and their decreasing heritability with age. These concepts were not followed up by systematic research, likely due to the difficulties involved in setting up large familial studies and perhaps the perceived limited clinical use of this kind of expensive and time-consuming project.

Detailed, high-granularity data on heritability by age are rare for most diseases. The familial heritability, clinical, and epidemiological statistics were available for eight prevalent LODs, Alzheimer’s disease (AD), type 2 diabetes (T2D), coronary artery disease (CAD), and cerebral stroke, and four late-onset cancers: breast, prostate, colorectal, and lung cancer. These statistics served as the basis for this study’s analysis and conclusions. This study investigated the model in which the polygenic risk of an individual remains constant with age and endeavored to establish how the higher odds of becoming ill of individuals with higher polygenic liability may lead to a change of risk allele distribution as the population ages and whether this alone may explain some of the known observational facts.

A set of computer simulations quantified the change in the risk allele representation for these LODs as the population ages and determined how and why these changes affect clinical predictive power and GWAS statistical discovery power with age more for some LODs than for others. Consequently, this study proposes a modification to GWAS cohort selection to improve statistical discovery power.

## Methods

### The model definition

According to Chatterjee et al. (2016), the conditional age-specific incidence rate of the disease, *I* (*t*|*G*) that is defined as the probability of developing the disease at a particular age *t*, given that a subject has been disease-free until that age, can be modeled using Cox’s proportional hazards model (Cox, 1972):

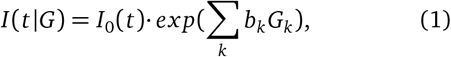

where *G* = (*G*_1_, &, *G*_*k*_) is the multiplicative effect of a set of risk factors on the baseline hazard of the disease *I*_0_(*t*). The set of age-independent variables in G could include genetic and environmental risk factors, as well as their interaction terms.

The following summary from Chatterjee et al. (2016) is particularly relevant to the methodology of this research: “logistic regression methods are preferred for the evaluation of multiplicative interactions. For case-control studies, if it can be assumed that environmental risk factors are independent of the SNPs in the underlying population, then case-only and related methods can be used to increase the power of tests for gene-environment interactions. To date, post-GWAS epidemiological studies of gene-environment interactions have generally reported multiplicative joint associations between low-penetrant SNPs and environmental risk factors, with only a few exceptions.” This means that the polygenic score *G* = Σ_*k*_ *b*_*k*_*G*_*k*_, as the lifelong characteristic of each individual, is used multiplicatively with *I*_0_(*t*), which encompasses environmental and aging effects.

It is important to note that the simulations conducted in this research rely on the model genetic architectures of the analyzed LODs, not a complete GWAS map of their experimentally discovered SNPs, because GWAS-discovered sets can explain only a fraction of these LODs’ heritability. These model genetic architecture SNPs are treated as “true” causal variants for disease liability and heritability, as discussed in Chatterjee et al. (2016), rather than GWAS-linked SNPs. They are used as *a priori* known constant causal SNPs that combine into individual polygenic risk scores (PRSs) for an LOD, as will be described further. The study by Pawitan et al. (2009), which followed the mathematical foundation and simulational validation of the liability model developed in Noh et al. (2006), served as a basis for the genetic architectures used in this study. Taking an aging population simulation approach allows for the identification of individuals becoming ill and, with them, the corresponding allele distribution between cases and controls, without intermediate steps and operating directly with the odds-ratio-based polygenic risk scores common to GWASs and clinical studies. The core of the simulation is Algorithm 1, operating on the known yearly incidence of an LOD and the PRSs for all individuals based on a modeled LOD genetic architecture:

#### Algorithm 1: Sampling individuals diagnosed with a disease proportionately to their polygenic odds ratio and incidence rate.

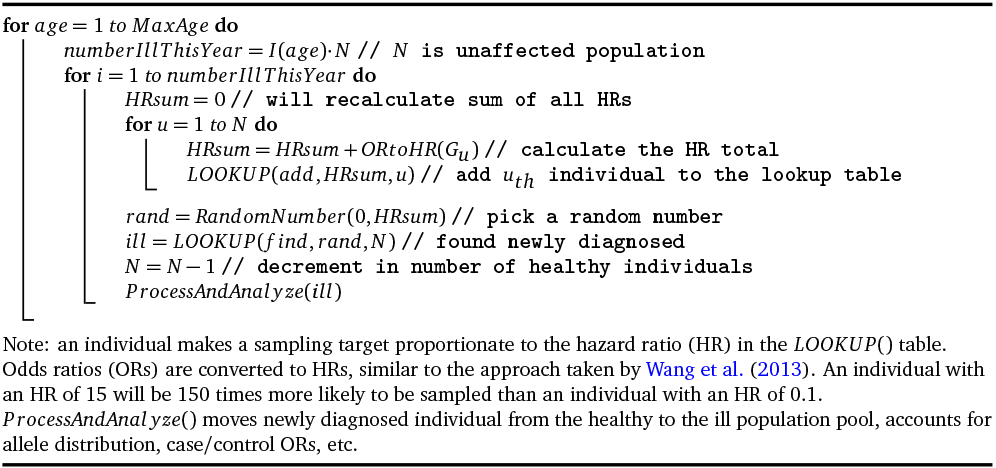

Descriptively, the algorithm works as follows. In this prospective simulation, each next individual to be diagnosed with an LOD is chosen proportionately to that individual’s relative PRS at birth relative to all other individuals in the as-yet-unaffected population. The number of individuals diagnosed annually is determined using the model incidence rate curve derived from clinical statistics. In this manner, the aging process is probabilistically reproduced using a population simulation model rather than a computational model. As the simulation progresses, the risk alleles are tracked for all newly diagnosed individuals and the remaining unaffected population, and their representation in the affected and remaining population is statistically analyzed.

The following sections describe the model genetic architectures, the LOD incidence models and the statistical foundations of this research.

### Allele distribution models

An in-depth review by Pawitan et al. (2009) extensively analyzed models of genetic architecture and through simulations determined the number of alleles required to achieve specific heritability and estimated the discovery power of GWASs. They calculated allele distributions and heritability and ran simulations for six combinations of effect sizes and minor allele frequencies (MAFs). Reliance on the conclusions of Pawitan et al. (2009) in this research makes it unnecessary to repeat the preliminary steps of evaluating the allele distributions needed to achieve the requisite heritability levels. The Pawitan et al. (2009) alleles represent the entire spectrum ranging from common, low-frequency, low-effect-size alleles to extremely rare, high-effect, high-frequency alleles. The five most relevant architectures were implemented in this study; see Table 1.

**Table 1.** Genetic architecture scenarios. Allele distributions as modeled by Pawitan et al. (2009).

| Scenario | MAF | Odds ratio |
| --- | --- | --- |
| A. Common low | 0.073–0.499 | 1.05–1.15 |
| B. Modest low | 0.0365–0.2495 | 1.05–1.15 |
| C. Rare low | 0.0146–0.0998 | 1.05–1.15 |
| D. Rare medium | 0.0146–0.0998 | 1.28–2.01 |
| E. Rare high | 0.0073–0.0499 | 1.63–4.05 |

It is also handy for repeatable allele tracking, rather than generating the continuous random spectrum of allele frequencies and effect sizes, to follow the Pawitan et al. (2009) configuration and discretize the MAFs into five equally spaced values within the defined range, with an equal proportion of each MAF and an equal proportion of odds ratios. For example, for scenario A, the MAFs are distributed in equal proportion at 0.073, 0.180, 0.286, 0.393, and 0.500, while the odds ratio (OR) values are 1.15, 1.125, 1.100, 1.075, and 1.05, resulting in 25 possible combinations. Having multiple well-defined alleles with the same parameters facilitated the tracking of their behaviors with age, LOD, and simulation incidence progression.

An individual polygenic risk score *β* can be calculated as the sum of the effect sizes of all alleles, which is by definition a log(OR) (natural logarithm of odds ratio) for each allele, also following Pawitan et al. (2009):

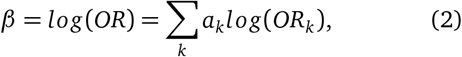

where *a*_*k*_ is the number of risk alleles (0, 1 or 2) and *OR*_*k*_ is the odds ratio of additional liability presented by the k-th allele.

Variance of the allele distribution is determined by:

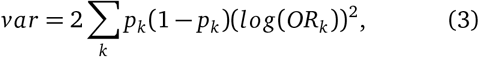

where *p*_*k*_ is the frequency of the k-th genotype (Pawitan et al., 2009).

The contribution of genetic variance to the risk of the disease is heritability:

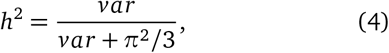

where *π*^2^/3 is the variance of the standard logistic distribution (Noh et al., 2006). For example, the number of variants needed for the Scenario A LODs is summarized in Table 2. Following Pawitan et al. (2009), the variants are assigned to individuals with frequencies proportionate to MAF *p*_*k*_ for SNP *k*, producing, in accordance with the Hardy–Weinberg principle, three genotypes (AA, AB or BB) for each SNP with frequencies 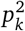, 2*p*_*k*_(1 − *p*_*k*_) and (1 − *p*_*k*_)^2^. The mean value *β*_*mean*_ of the population distribution can be calculated using the following equation:

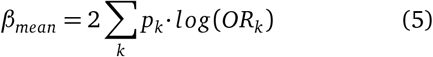

**Table 2.**
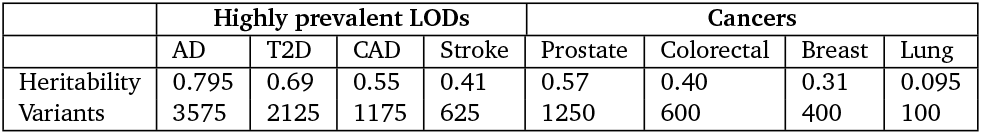
Heritability of analyzed LODs and an example of required variant numbers for common low-effect variants: Scenario A.

|  | Highly prevalent LODs |  |  |  | Cancers |  |  |  |
| --- | --- | --- | --- | --- | --- | --- | --- | --- |
|  | AD | T2D | CAD | Stroke | Prostate | Colorectal | Breast | Lung |
| Heritability | 0.795 | 0.69 | 0.55 | 0.41 | 0.57 | 0.40 | 0.31 | 0.095 |
| Variants | 3575 | 2125 | 1175 | 625 | 1250 | 600 | 400 | 100 |

Customarily, the individual PRSs are normalized relative to *G*_*mean*_, resulting in a zero mean initial population PRS, making it easy to compare higher- and lower-risk individuals. The figures S13 and S14 in Supplementary Chapters depict the corresponding population distribution of detrimental variants and PRSs for the common low-effect-size genetic architecture.

A part of simulation functionality is to allocate the genetic architectures and calculate the variance, using Eq. (3), of each genetic architecture instance described above. Each genetic architecture listing is represented in the Supplementary Data executable folder; for example, the file “CommonLow.txt” lists the variants describing Scenario A (only three columns are used for this simulation: SNP—-internal use identifier, EAF—-effect allele frequency, and OR). In the case of Scenario A, *var* = 0.09098 for the single set of SNPs listed in this file. Rearranging Eq. (4) and changing the multiple allows for the discovery of the number of variant sets for each LOD, as seen in Table 2, and for the target heritability to be closely approximated. Each simulation run calculates the PRS variance within the population and records heritability and allele distributions for the case and control populations as the simulated age progresses.

### Evaluating GWAS statistical power

GWAS statistical power is the estimate of the ability of GWASs to detect associations between DNA variants and a given trait, and depends on the experimental sample size, the distribution of effect sizes, and the frequency of these variants in the population (Visscher et al., 2017). Statistical power calculations are very useful in a case/control study design for determining the minimum number of samples that will achieve adequate statistical power; conventionally, statistical power of 80% is considered to be acceptable (Hong and Park, 2012). To achieve greater power, a disproportionately larger number of cases and controls may be required, which is frequently unrealistic for cohort studies. A number of statistical power calculators are available, for example, Sham and Purcell (2014). This study utilized the Online Sample Size Estimator (OSSE) (Online Sample Size Estimator).

The progress made by GWASs over the last decade, particularly in relation to polygenic traits, was to a large extent due to ever-increasing cohort sizes. Cohort size is one of the principal factors limiting GWAS discovery power, making it an important benchmark for this study. Here, the cohort size is defined as the number of cases and controls needed to achieve 80% statistical discovery power when the case/control allele frequency changes with cohort age for a subset of representative alleles in the model genetic architectures. For each such allele in the simulated population, the allele frequency for cases and controls is tracked as age progresses. The difference between these MAFs gives the non-centrality parameter (NCP) *λ* for two genetic groups (Sham and Purcell, 2014; Luan et al., 2001):

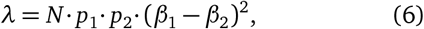

where N is the overall population sample size and *p*_1_ *andp*_2_ the fractions of cases and controls, and *β*_1_ and *β*_2_ are the case and control mean log(OR) for an allele of interest. The values *p*_1_ = *p*_2_ = 0.5, or an equal number of cases and controls, are used throughout this publication. Having obtained NCP *λ* from Eq. (6), Luan et al. (2001) recommended using SAS or similar statistical software to calculate the statistical power, using the following SAS statement:

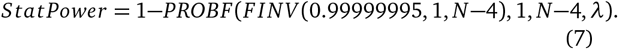

The conversion of this equation to its R equivalent, which was used to process the simulation output, is:

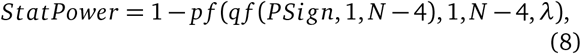

where *PSign* = 0.99999995 corresponds to a 5 · 10^−8^ significance level. The outputs of this conversion were validated using the Online Sample Size Estimator (OSSE) (Online Sample Size Estimator). This equation returns statistical power based on a case/control number and the NCP as calculated above. To find the number of cases needed for 80% GWAS discovery power, having the (*β*_1_ − *β*_2_), a rapid convergence R routine was used to iterate the values of *N* until the value of *Stat Power* matched 0.8 (80%) with an accuracy better than ±0.01% for each age and allele distribution of interest.

### LOD incidence functional approximation

Chapter S3 in Supplementary Chapters describes the functional approximations of the yearly incidence of Alzheimer’s disease, type 2 diabetes, coronary artery disease, and cerebral stroke, and four late-onset cancers: breast, prostate, colorectal, and lung cancer. As a short summary, for all of the above LODs, the incidence rate curves can be approximated during the initial disease onset periods with an annual incidence growth that is close to exponential. This exponential growth continues for decades; see Table 3 and Chapter S3 in Supplementary Chapters.

**Table 3.** Age to which LOD incidence rate rises exponentially.

|  | Highly prevalent LODs |  |  |  | Cancers |  |  |  |
| --- | --- | --- | --- | --- | --- | --- | --- | --- |
|  | AD | T2D | CAD | Stroke | Prostate | Colorectal | Breast | Lung |
| Age (years) | 103 | 55 | 81 | 79 | 48 | 62 | 72 | 70 |

Later, the growth may flatten in old age, as is the case with T2D, slow down, as is the case with cerebral stroke and CAD, or continue exponentially to a very advanced age, as is the case with Alzheimer’s disease. An R script automates the determination of the best fit for logistic and exponential approximation from the clinical incidence data.

### Sampling based on the LOD incidence rate and individual PRS

The incidence rate functional approximations for the analyzed LODs are used to find the average number of diagnosed individuals *N*_*d*_ for each year of age *t* as a function of the incidence rate *I* (*t*) and the remaining population unaffected by the LOD *N*_*u*_(*t*) in question:

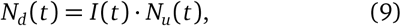

In the next year of age, the unaffected population will have been reduced by the number of individuals diagnosed in the previous year *N*_*d*_(*t*):

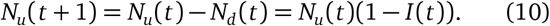

The number of individuals projected to become ill per year, as well as the remaining unaffected population, is applied in Algorithm 1.

For the PRS of the simulated population based on odds ratios built using (Pawitan et al., 2009) model, if an LOD is characterized by low incidence within an age interval, and the odds ratio is close to 1, odds ratio values are practically identical to hazard ratio or relative risk (RR). For example, Song et al. (2014) treat OR and RR as equivalent in the case of breast cancer in their simulation study. For higher values, an OR usually significantly exceeds the RR. An adjustment formula by Zhang and Kai (1998) can provide OR to hazard ratio approximation.

### Individual values analysis and cohort simulation

It can be expected that, for an LOD with higher incidence and heritability, the fraction of the highest-PRS individuals will diminish more rapidly with age. For such LODs, the relatively-lower-PRS individuals will represent the bulk of the LOD cases at an earlier age compared to LODs with lower incidence and heritability. The LODs are characterized by a wide range of heritability and progression patterns of incidence rate with age. For example, T2D and breast cancer begin their incidence rise relatively early but reach quite different levels at older age, while colon cancer and AD start later and also reach quite different maximum incidence and cumulative incidence levels; see Fig. S1 in Supplementary Chapters. In the absence of mortality, both due to general frailty and other LODs, the incidence progression makes it appear as though, sooner or later, depending on the incidence magnitude, the majority of the population would be diagnosed with every LOD. In reality, this does not happen because of ongoing mortality from all causes.

Two main LOD simulation types are described next:

1. **The individual values analysis of polygenic risk scores and risk allele frequency for individuals diagnosed with a disease at each specific age and the remaining population at this age.** The abbreviation “IVA” is used interchangeably with “individual values analysis” in this publication. The IVA uses one-year age slices and is performed as follows. Initially, the mean and variance of the PRS for the whole population are calculated. Next, based on the required incidence value for each year, individuals are picked from the unaffected population by randomly sampling the population with a probability proportionate to the individual’s PRS, as summarized in Algorithm 1. These individuals become the cases for the relevant year’s IVA, and the mean and variance of their PRSs are also calculated and recorded. Mortality does not need to be applied to the IVA scenario because it affects the future cases and controls in equal numbers, and accounting for mortality would only result in a smaller population being available for analysis. To track the GWAS statistical discovery power, the same nine representative variants (configurable) are tracked for all LODs simulated. The process continues in this way until the maximum desired simulation age is reached.
2. **A simulated cohort study for each of these diseases.** For the sake of brevity, the word “cohort” is also used throughout this publication. The clinical study cohort simulation performs an analysis identical to that described above. The difference is that, here, the simulated GWAS clinical studies are performed with a patient age span of 10 years, which is a typical cohort age span, although any age span can be chosen as a simulation parameter. The simulation statistics are collected using the mid-cohort age, which is the arithmetic half-age of the cohort age span. In the first simulation year, a population equal to one-tenth of the complete population goes through the steps described for IVA. Each year, an additional one-tenth starts at age 0, while the previously added individuals age by one year. This continues until all 10 ages are represented. This combined cohort proceeds to age and is subject to the disease incidence rate and mortality according to each individual’s age.

Mortality is applied, with a probability appropriate to each year of age, to both accumulated cases and controls. As the population ages, both the case and control pool numbers diminish. Take, for example, a cohort study that includes a 10-year span, say, between 50 and 59 years of age. The cases for the cohort are composed of individuals who were diagnosed with an LOD at any age either younger than or including their current age, producing a cumulative disease incidence over all preceding years of age. For example, some of the individuals that are cases now, at age 59, may have been healthy at age 58. Some of the controls in our cohort at the age of 51 may or may not be diagnosed at an older age, which would qualify them as cases for this cohort, but they are currently younger and healthy. Therefore, it can not be known with certainty the extent to which younger controls differ from cases, except for the fact that they are not currently diagnosed—not unlike a real statistical study cohort. As a result, the corresponding GWAS discovery power can be expected not to change as dramatically as it does for the individual values analysis.

The following additional mortality scenarios were also simulated: (a) double mortality for cases compared with the unaffected population, (b) no mortality for either cases or controls, and (c) a one-year age span cohort with no mortality for either cases or controls.

The youngest age cohort for each LOD is defined as the mid-cohort age at which the cumulative incidence for a cohort first reaches 0.25% of the population. For consistency, this threshold was considered in this study as the minimum cumulative incidence age, allowing for the formation of well-powered cohort studies for all analyzed LODs.

### The simulation design summary

Preliminary data collection and analysis steps are shared by all simulation runs and include: (a) preparing the genetic architecture files and calculating the number of variants needed, based on each modeled LOD heritability, as described in the “Allele distribution models” section above, and (b) determining the parameters of functional approximation for LOD incidence from published statistics, as described in Chapter S3 in Supplementary Chapters.

A simulation run for a single LOD can be logically divided into the following four steps:

1. Build the gene variants pool as outlined in the “Allele distribution models” section and load the incidence rate functional approximation parameters.
2. Allocate population objects and assign individual PRSs. Allocate all other simulation objects and arrays that will be used by the simulation.
3. Run the simulation’s Algorithm 1 from age 0 to 100 for either the IVA or the cohort study scenario, described in the “Individual values analysis and cohort simulation” section. Calculate and record the simulation data in comma separated value files.
4. Determine statistical power for cases and controls for each cohort based on the “Evaluating GWAS statistical power” section.

The above steps were completely reinitialized and performed for each LOD analysis. The complete simulation iterated through all eight LOD analyses in two scenarios: a per-year-of-age population IVA and a simulated GWAS cohort study.

### Validation simulations

Based on the model described above, it can be expected that the allele distribution in a population of the same age with a given initial genetic architecture will depend solely on cumulative incidence, which represents the fraction of the population that succumbs to a disease. The purpose of validation simulation runs performed with (a) constant, (b) linear and (c) exponential incidence rates was to validate whether or to what extent this expectation is correct and whether the outcomes would differ between various genetic architectures. The validation simulations confirmed that PRSs for the population controls and cases, viewed in the individual values analysis at every age, depend on the cumulative incidence and the LOD heritability, and are independent from the incidence progression shape within each genetic architecture. The procedures used in the validation simulations are described in Chapter S2 in Supplementary Chapters.

### GWAS simulations and effect size adjustment for younger cases and older controls cohorts

The case-control populations produced by the above simulations were suitable for consequential simulated GWAS association analysis. The simulations saved the output in PLINK format (Chang et al., 2015; Purcell and Chang, 2019). The initial validation, analysis and file format conversions were performed using PLINK v1.9. The logistic regression GWAS with adjustment for age was performed using R script *AdjustByAge.R*, as described below, and the outputs were validated with Regression Modeling Strategies (rms) GWAS R package by Harrell Jr et al. (2018) and PLINK, both achieving individual SNP association results identical to produced by this R script.

The GWAS simulations showed that the apparent effect size tended to increase with age of control cohort, when analyzed against the youngest possible cohort, in comparison to a “true” value, which was chosen as the effect size value from the youngest age-matched cohort. Chatterjee et al. (2003) demonstrated application of age bias to a leprosy case-control study, using *logit P* = *β*_0_ + *β*_1_*T* + *β*_2_*S*, where the bias function T is expressed as *T* = 100(*Age* + 7.5)^−2^ in that study.

The R script *AdjustByAge.R*, based on R Generalized Linear Models glm() functionality (R Core Team, 2013), performed the GWAS association and iterative age covariate adjustment starting with a youngest possible age-matched cohort, and proceeding with the progressively older control cohorts. The best match bias adjustment functions and their parameters differed between LODs. The simulation results showed that the differential increase in the value of the effect size was approximately proportionate to the effect size magnitude for all analyzed here LODs. The differential normalized effect size *D*(*t*) can be expressed as:

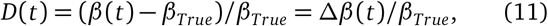

where *β* (*t*) and *β*_*True*_ are the effect size value found for older control cohort compared to a known “true” effect size. The variable *D*(*t*) will be referred further as normalized bias. The GWAS simulations associated the effect sizes in 5 year control cohort age increments, and matched the best power exponent regression function:

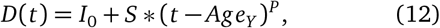

where *I*_0_ ans *S* are the linear regression intercept and slope, *t* is an older control cohort age, and *Age*_*Y*_ is the young case cohort age, and *P* is the best match power exponent. When the solution to Eq. (12) is correctly estimated for one gene variant (likely for an SNP with a larger effect size), it could be used to adjust other discovered variants’ effect sizes using rearranged Eq. (11) and Eq. (12). The R script *FindAdjustmentRegressionFunction.R* implementing lm() linear regression iteratively fitted the best matching adjustment power with very lowest residuals. Additionally this script evaluated the regression with fixed P=2 (quadratic regression) for all LODs. The data preparation and scripting steps described here are listed with specifics in *GwasSimulationPipeline.txt*, available along with the R scripts in Supplementary Data.

### Data sources, programming and equipment

The population mortality statistics from the US Social Security Actuarial Life Table provided yearly death probability and survivor numbers up to 119 years of age for both men and women.

Disease incidence data from the following sources were extensively used for analysis, using the materials referenced in Chapter S1 in Supplementary Chapters for corroboration: Alzheimer’s disease: (Brookmeyer et al., 1998; Edland et al., 2002; Kokmen et al., 1988; Hebert et al., 1995); type 2 diabetes: (Boehme et al., 2015); coronary artery disease and cerebral stroke: (Rothwell et al., 2005); and cancers: (Cancer Statistics for the UK; Kuchenbaecker et al., 2017).

The simulations were performed on an Intel i9-7900X CPU-based 10-core computer system with 64GB of RAM and an Intel Xeon Gold 6154 CPU-based 36-core computer system with 288GB of RAM. The simulation is written in C++ and can be found in Supplementary Data. The simulations used population pools of 2 billion individuals for the LOD simulations and 300 million for validation simulations, resulting in minimal variability in the results between runs.

The cohort simulations were built sampling at least 5 million cases and 5 million controls from the surviving portion of the initial 2 billion simulated individuals, which is equivalent to 0.25% of the initial population. This means that the cohort study would begin its analysis only when this cumulative incidence was reached. Conversely, the analysis would cease when, due to mortality, the number of available cases or controls declined below this threshold. For all LODs, this maximum mid-cohort age was at least 100 years and, depending on LOD, up to a few years higher. This confirms that, as described later in the Discussion section, in cohorts composed of younger cases and older controls it is feasible to form control cohorts up to 100 years of age.

The simulation runs for either all validation scenarios or for a single scenario for all eight LODs took between 12 and 24 hours to complete. The final simulation data, additional plots and elucidation, source code, and the Windows executable are available in Supporting Information. Intel Parallel Studio XE was used for multi-threading support and Boost C++ library for faster statistical functions; the executable may be built and can function without these two libraries, with a corresponding slowdown in execution. The ongoing simulation results were saved in comma separated files and further processed with R scripts during subsequent analysis, also available in Supplementary Data.

### Statistical analysis

Large variations between simulation runs complicate the analysis of population and genome models. This issue was addressed in this study by using a large test population, resulting in negligible variability between runs. The statistical power estimates deviated less than 1% in a two-sigma (95%) confidence interval, except for the early Alzheimer’s disease cohort, which commenced at 1.5% and fell below the 1% threshold within 4 years (see \**TwoSDFraction.csv* files in Supplementary Data). In addition to ensuring that the simulations operated with reliable data, this eliminated the need for the confidence intervals in the graphical display.

The GWAS simulations and variant effect size covariate adjustment by age were more memory intensive and the 200 million simulated population with 500 thousand case and control cohorts was possible with the above equipment. In this instance, two-sigma confidence intervals for simulated GWAS discovery and regression parameters are presented in the corresponding plots.

## Results

### Validation simulations for all genetic architectures

The validation simulations for all scenarios described in Methods Table 1 were performed not as models of specific diseases but to determine the behavior of all allele scenarios and the resulting allele frequency changes under simple controlled and comparable-to-each-other incidence scenarios. It was important to characterize all genetic architectures and to identify the differences and similarities in behavior between them.

These simulations confirmed that a change in the population’s mean PRS and a change in the cases’ mean PRS, viewed as instantaneous values for each age, are dependent on the cumulative incidence and the magnitude of the initial genetic model heritability. If mortality is excluded, they are not dependent on the shape of incidence progression with age (see Fig. S2 in Supplementary Chapters) and are qualitatively similar between the genetic architectures (see Fig. S3 in Supplementary Chapters). This means that, when the same level of cumulative incidence is reached with any incidence pattern, the allele distribution for the diagnosed cases and the remaining unaffected population is identical.

### Analysis of common, low-effect-size genetic architecture scenario

The simulation results for the eight analyzed LODs are presented next.

The IVA and cohort simulations were performed for all genetic architecture scenarios, from low to high effect sizes, and common to low allele frequencies, and the results were found to be qualitatively consistent between all these scenarios. As a consequence, this report primarily focuses on the common low-effect-size genetic architecture scenario A, which the latest scientific consensus considers to be the genetic architecture behind the majority of polygenic LODs; the results are virtually identical for scenarios B and C, as validated in Fig. S3 in Supplementary Chapters, making it unnecessary to present separate figures for these two scenarios.

The scatter plots in Fig. 1 show the distributions of PRS for cases diagnosed as age progresses for the common, low-effect-size genetic architecture scenario A. The PRSs of individuals diagnosed with an LOD and the age-related change of the average LOD PRS of the unaffected population are demonstrated in Fig. 2. The color bands show a one standard deviation spread for cases and controls, which, in the case of newly diagnosed cases, represents approximately two-thirds of the diagnoses at each age. This figure demonstrates how the initially high average polygenic risk of newly diagnosed cases declines as the most predisposed individuals are diagnosed with each passing year of age. The average PRS of the unaffected population decreases much more slowly. At advanced old age, the average polygenic risk of the newly diagnosed is lower than the risk for an average individual in the population at a young age; this is true for all four highly prevalent LODs: AD, T2D, CAD, and stroke.

**Figure 1.**
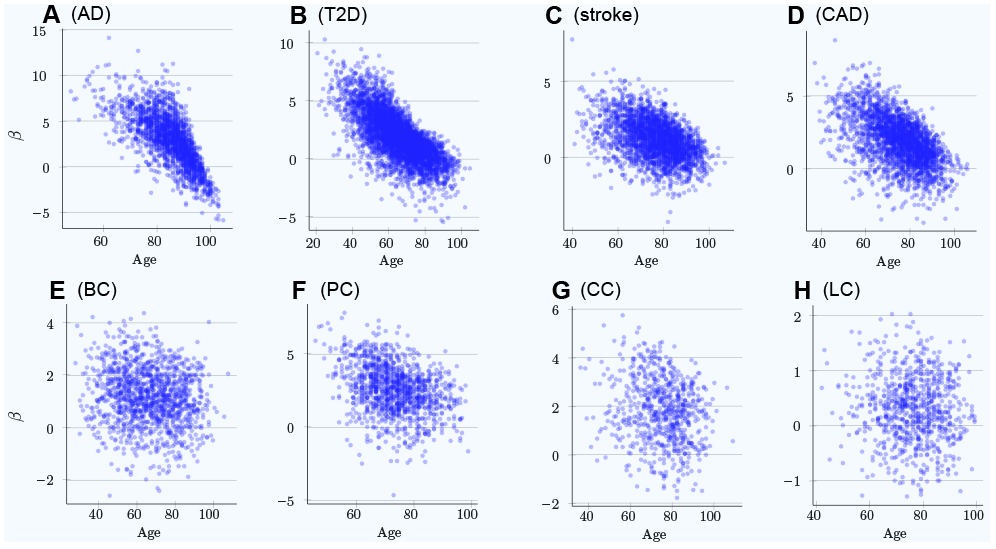
Polygenic risk scores of individuals diagnosed with an LOD as a function of age. **(A)** Alzheimer’s disease, **(B)** type 2 diabetes, **(C)** cerebral stroke, **(D)** coronary artery disease, **(E)** breast cancer, **(F)** prostate cancer, **(G)** colorectal cancer, **(H)** lung cancer. Scatter plots show the distributions of PRS for cases diagnosed as age progresses, with ongoing mortality. *β* = *log*(*OddsRatio*) visually implies the underlying heritability and incidence magnitudes. If the regression line can be easily drawn, dropping diagonally as age progresses, there is a combination of high heritability and increasing cumulative incidence. Otherwise, a plot appears as a relatively symmetrical blob.

**Figure 2.**
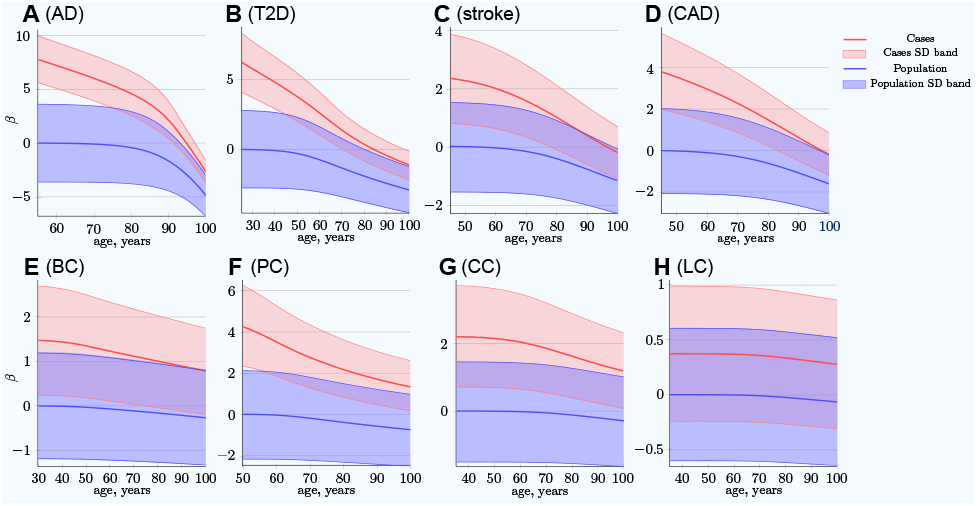
Polygenic risk score difference between newly diagnosed individuals and the remaining unaffected population. **(A)** Alzheimer’s disease, **(B)** type 2 diabetes, **(C)** cerebral stroke, **(D)** coronary artery disease, **(E)** breast cancer, **(F)** prostate cancer, **(G)** colorectal cancer, **(H)** lung cancer. *SD band* is a band of one standard deviation above and below the cases and the unaffected population of the same age. For highly prevalent LODs, at very old age, the mean polygenic risk of new cases crosses below the risk of an average healthy person at early onset age. *(Common low-effect-size alleles (scenario A), showing largest*-*effect variant with MAF* = *0.5, OR* = *1.15)*.

This phenomenon is a consequence of the effect allele frequency change, in which the highest-effect alleles show the greatest difference between the diagnosed and the remaining unaffected population as well as the fastest change in frequency difference with age. Statistically, individuals possessing the higher-risk alleles are more likely to succumb and to be diagnosed earlier, thus removing the allele-representative individuals from the unaffected population pool; see Fig. 3. These plots show the most dramatic change for AD and T2D—the LODs with the highest cumulative incidence and heritability. The smallest change corresponds to the LOD with the lowest incidence and heritability: lung cancer.

**Figure 3.**
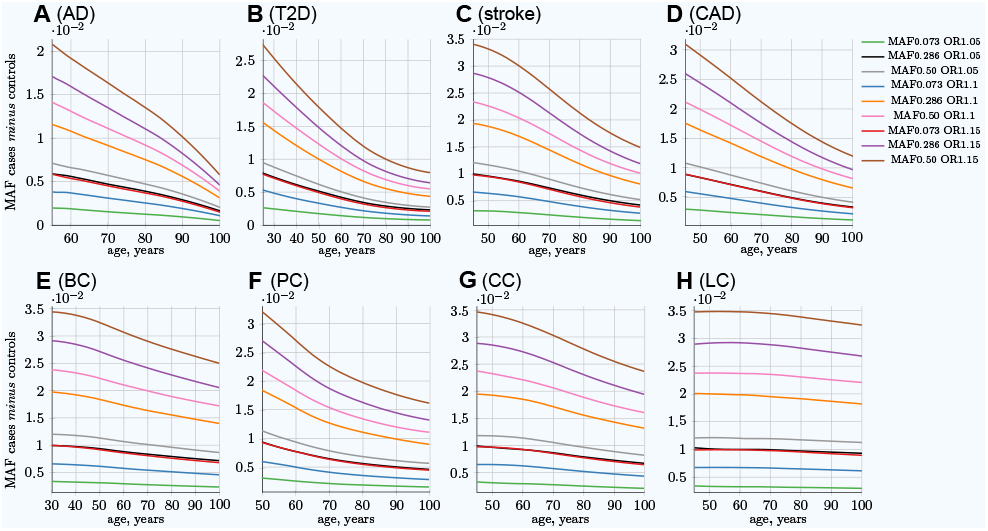
Allele frequency difference between newly diagnosed individuals and remaining population of the same age. **(A)** Alzheimer’s disease, **(B)** type 2 diabetes, **(C)** cerebral stroke, **(D)** coronary artery disease, **(E)** breast cancer, **(F)** prostate cancer, **(G)** colorectal cancer, **(H)** lung cancer. The **MAF cases** *minus* **controls** value is used to determine GWAS statistical power; see Eq. (6). Rarer and lower-effect-size (OR) alleles are characterized by a lower MAF relative change, see also Fig. S5 in Supplementary Chapters. *(Displayed here: 9 out of 25 SNPs, which are described in Methods for common low*-*effect*-*size alleles* - *scenario A)*.

It is important to note that the absolute MAF values for cases diagnosed at a particular age and controls do not change much with age progression for all LODs. For example, for T2D, the allele frequency for the allele with an OR of 1.15 and an initial population MAF of 0.2860 is 0.2860 for controls and 0.3088 for cases at the age of 25. This changes to 0.2789 for controls and 0.2871 for cases at the age of 80 in the IVA case—a change of only a few percentage points. At the same time, the relative differences change correspondingly from 0.0228 to 0.0081, a 2.7 times change, which is very significant for GWAS discovery power, as can be seen in Eq. (6). The absolute MAF change is even less prominent in the cohort scenario, as can be seen in Fig. S13 in Supplementary Chapters, which shows the same allele. The small change in the absolute value for older age groups makes it difficult to analyze this effect using, for example, GWAS SNP database statistics for different age groups. The effect would be hidden behind interpersonal and populational genetic variability in hundreds and thousands of SNPs, changing their balance slightly with age in the case of the common, low-effect-size genetic scenario. This effect is long established for highly detrimental variants such as the BRCA1/2 gene mutations in the case of breast cancer (Kuchenbaecker et al., 2017) and the APOE e4 allele in the case of AD (Farrer et al., 1997), where these gene variant carriers are known to be present in lower numbers among older undiagnosed individuals.

The cohort simulation shows a much more averaged change for these same scenarios because cohorts represent accumulative disease diagnoses from earlier ages, while mortality removes older individuals; see Fig. S5 in Supplementary Chapters. While the MAF difference between cases and controls shown in the above figures is illustrative by itself, it is most important for determining the GWAS statistical discovery power using Eq. (6) and Eq. (8) and from there the number of cases necessary to achieve 0.8 (80%) statistical power. From these equations, it is apparent that GWAS statistical discovery power diminishes as a complex function of a square of case/control allele frequency difference. The age-related change in the number of cases needed to achieve 80% GWAS discovery power for an age-matched case/control cohort study is presented in Fig. 4.

**Figure 4.**
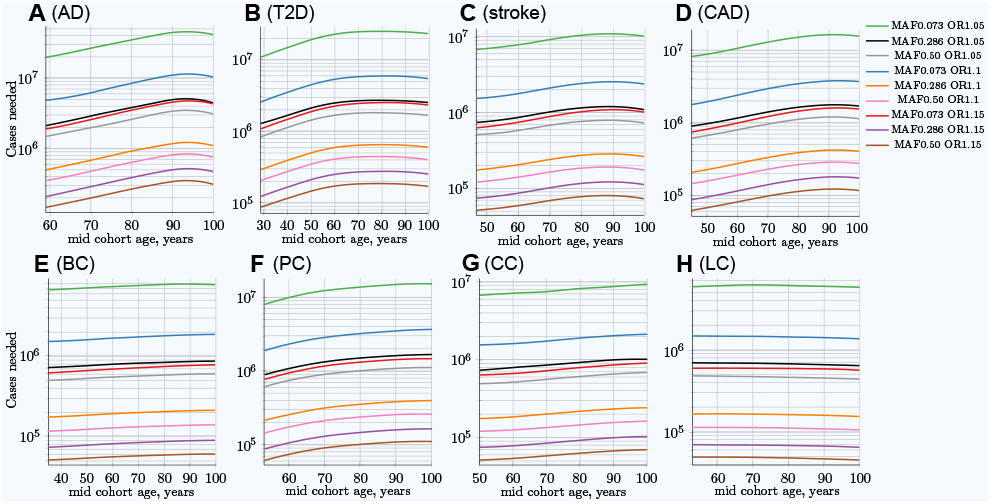
Change in number of cases needed to achieve 80% discovery power in age-matched cases and controls cohort design. **(A)** Alzheimer’s disease, **(B)** type 2 diabetes, **(C)** cerebral stroke, **(D)** coronary artery disease, **(E)**breast cancer, **(F)** prostate cancer, **(G)** colorectal cancer, **(H)** lung cancer. Age-matched cohorts require larger numbers of participants to achieve the same GWAS discovery power compared to the youngest cohort age. There is a noticeable difference between cancers (with the exception of prostate cancer; see the Discussion section) and other LODs. *(Displayed here: 9 out of 25 SNPs, which are described in the Methods section for common, low*-*effect*-*size alleles*— *scenario A)*.

In the hypothetical IVA case, the number of individuals required to achieve the desired GWAS discovery power increases rapidly; see Fig. S6 in Supplementary Chapters. This is a quite informative instantaneous value of statistical power; however, neither GWASs nor clinical studies ever consist of individuals of the same age, due to the need to have a large number of individuals to maximize this same statistical power. The cohort scenario is correspondingly less extreme, as seen in Fig. 4. These plots show an increase in the number of participants needed to achieve adequate GWAS statistical power between the lowest effect and frequency and the highest effect and frequency alleles; this number exhibits a greater-than-hundredfold variation between alleles within the genetic architecture. The required number of cohort participants is quite similar for the same effect alleles among all eight LODs; for example, the highest-effect allele for each LOD requires 5· 10^4^−1.4· 10^5^ cases for 80% GWAS discovery power at younger ages. The change in allele frequency with age between cases and controls shows substantial variation among LODs, with the greatest change occurring in AD and the least significant in lung cancer; see Fig. 3. These cohort results are simulated with identical mortality for cases and controls. Mortality has an impact on the cohort allele distribution.

Table 4 combines the heritability and incidence of the LODs with the summarized simulation results from the cohort simulation, also seen in Fig. 4 and Fig. S4 in Supplementary Chapters.

**Table 4.**
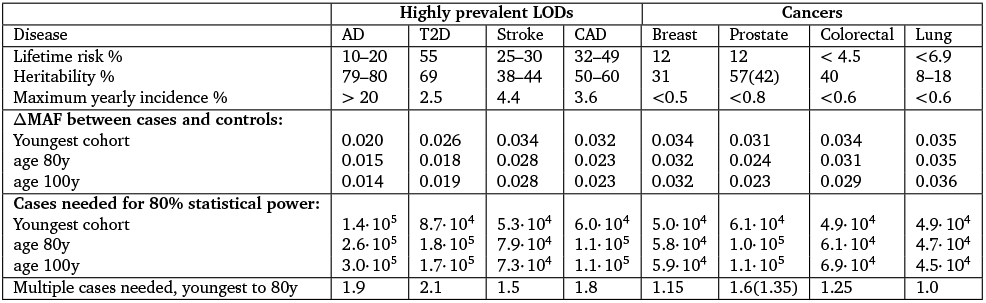
LOD statistics and age-matched cohort simulation summary. The MAF values and cases needed for 0.8 (80%) GWAS statistical discovery power are for the common, low-effect-size alleles, scenario A. Cohorts span 10 years. The results shown are for the allele with a MAF of 0.5 and an OR of 1.15, the largest effect allele, which requires the smallest number of cases/controls. “Maximum incidence %” is the largest incidence at older age. “Case mult.” is the multiple of the number of cases needed for the 80-year-old cohort to achieve the same statistical power as the early cohort. Prostate cancer heritability is 57%, according to Hjelmborg et al. (2014). Shown in braces, 42% heritability (Grönberg, 2003), which is more in line with the other three cancers.

### Validating more extreme mortality scenarios

More extreme mortality scenarios—both lower and higher—than one could expect in a real cohort study were validated in this set of simulations. The results were relatively close to those presented for equal case/control mortality. The extreme cases of (a) no mortality for either cases or controls and (b) double the mortality of cases compared to controls produce very similar allele distributions before the age of 85, while diverging somewhat at older ages. The scenario in which the mortality of cases is double that of controls is higher than the clinically known mortality for the analyzed LODs. While this may have been a realistic scenario a century ago, before modern healthcare, it is certain that patient mortality is lower these days. In addition, a one-year cohort without mortality was used as the most extreme validation case. This scenario can also be considered an individual cumulative case, which counts everyone who became ill by a specified age as cases and everyone healthy at that age as controls. These validation cohort scenarios are summarized in Fig. S7 in Supplementary Chapters.

The mortality analysis was applied to one LOD at a time. This research did not attempt to estimate increased mortality for multiple disease diagnoses. Collerton et al. (2009) followed a cohort of individuals over the age of 85 in Newcastle, England, and found that, out of the 18 common old-age diseases they tracked, a man was on average diagnosed with four and a woman with five, not to mention a plethora of other less common diseases and their causal share in individual mortality.

### Evaluating rare, medium-effect-size genetic architecture scenario

Other genetic architecture scenarios produce qualitatively similar patterns, specifically differing in the number of cases needed to achieve 80% statistical power for medium- and large-effect genetic architecture scenarios. The rare, medium-effect-size allele (scenario D) results are presented in Fig. S8, Fig. S9, Fig. S10, and Fig. S11 in Supplementary Chapters. There, at younger ages, the MAF difference between cases and controls is larger for rare, medium-effect-size alleles. The number of cases and controls needed to achieve 80% GWAS statistical power for all eight LODs is approximately five times lower, a direct consequence of these variants’ larger effect sizes. This result perhaps excludes the scenario of rare, medium-effect-size alleles being causally associated with the LODs reviewed here, because GWAS studies would be more readily able to discover a large number of causal SNPs. From a qualitative perspective, all reviewed genetic architecture scenarios provide similar patterns of increasing numbers of cohort study cases needed to maintain the same discovery power with age progression.

### Advantage of using youngest possible case cohorts and oldest control cohorts

The scenarios simulating the number of cases needed when the case cohort uses the youngest possible participants with increasingly older control cohorts are presented in Fig. 5 and Fig. S12 in Supplementary Chapters. The multiplier representing the decrease in the number of cases that are needed in this scenario is represented by the blue lines in Fig. 6, which strongly contrasts with the number of cases needed for the same GWAS discovery power in a classic age-matched study design demonstrated in the red line in Fig. 6, which increases with age. The summary of the comparison of the two cohort designs is presented in Fig. 7.

**Figure 5.**
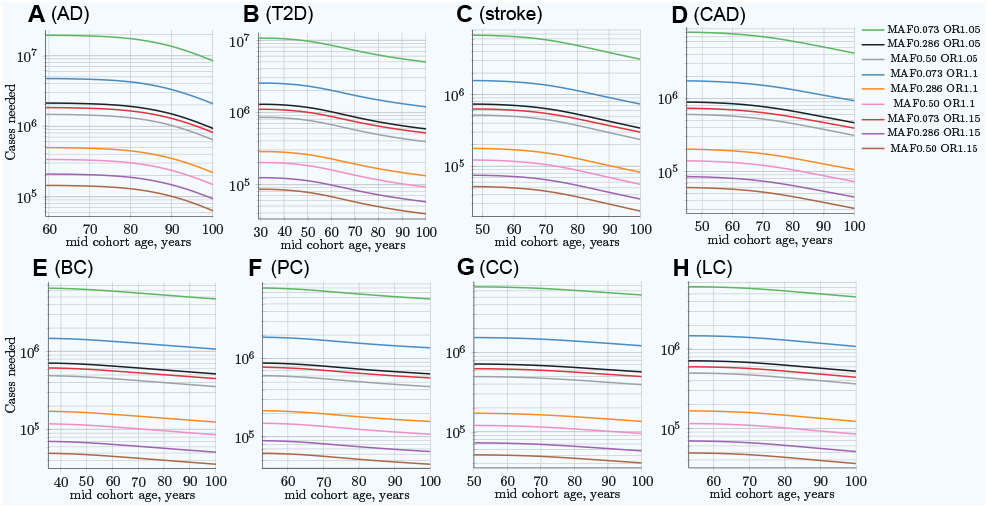
Change in number of cases needed for 80% discovery power in a cohort study when using progressively older controls compared to fixed-age young cases. **(A)** Alzheimer’s disease, **(B)** type 2 diabetes, **(C)** cerebral stroke, **(D)** coronary artery disease, **(E)** breast cancer, **(F)** prostate cancer, **(G)**colorectal cancer, **(H)** lung cancer. Cases’ mid-cohort age is leftmost age (youngest plot point); control mid-cohort ages are incremental ages. The number of cases needed for 80% discovery power is smaller when using older controls, particularly for those LODs showing the most prominent increase in the number of cases needed for older age in matched-age cohorts, as can be seen in Fig. 4. *(Displayed here: 9 out of 25 SNPs, which are described in the Methods section for common, low*-*effect*-*size alleles*—- *scenario A)*.

**Figure 6.**
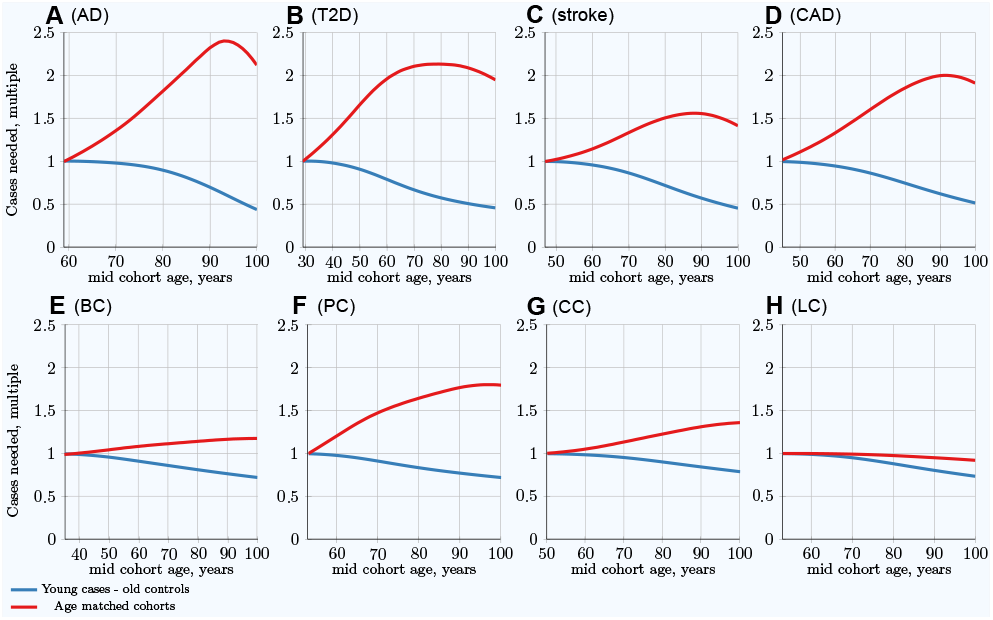
Per LOD comparison: Youngest possible cases and increasingly older controls *vs* classical age-matched cohorts. **(A)** Alzheimer’s disease, **(B)** type 2 diabetes, **(C)** cerebral stroke, **(D)** coronary artery disease, **(E)** breast cancer, **(F)** prostate cancer, **(G)** colorectal cancer, **(H)** lung cancer. The multiplier showing the reduction in the number of cases needed in a young cases–older controls scenario is shown in blue, strongly contrasting with the number of cases needed for the same GWAS discovery power in a classic age-matched study design, shown in red, which increases with age. *(Common low*-*effect*-*size alleles scenario A.)*

**Figure 7.**
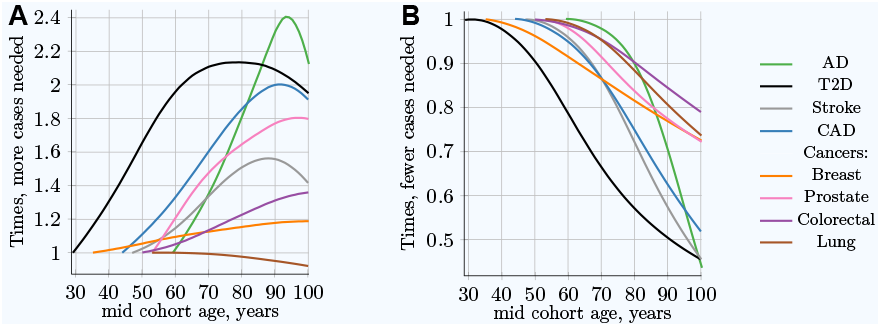
Advantage of using youngest possible cases and increasingly older controls, compared to classical age-matched cohorts. **(A)** Relative **increase in number of cases needed** for 80% discovery power in a cohort study using progressively older **case and control cohorts of the same age**. **(B)** Relative **decrease in the number of cases needed** for 80% discovery power in a cohort study when using **progressively older control cohorts compared to fixed-age young-case cohorts**. The youngest age cohort for each LOD is defined as the mid-cohort age at which the cumulative incidence for a cohort first reaches 0.25% of the population. Therefore, the leftmost point on each LOD line is the reference (youngest) cohort, and as cohorts age, the cohort case number multiple required to achieve 0.8 statistical power is relative to this earliest cohort. While all alleles display a different magnitude of cases needed to achieve the required statistical power, the change in the multiplier with age is almost identical for all alleles within a given genetic architecture scenario. *(Common low*-*effect*-*size alleles scenario A.)*

Thus, cohorts composed of the youngest possible cases and the oldest available controls improve the discovery power of GWASs. Equivalently, such cohorts require a smaller number of participants to achieve the same discovery power.

### Characterizing and adjusting for effect size in the younger cases and older controls GWAS

The GWAS association simulations with the youngest possible cases and older controls cohorts showed the SNP effect size exceeding the known “true” effect size with the increasing controls cohort age. This is the expected consequence of the larger effect allele differential between these cohorts compared to the age matched cohorts. For SNPs defined in the simulation with the effect size 0.14 (OR=1.15), the association analysis found effect sizes near 0.20 (OR=1.21) for CAD, and stroke with 100 years old control cohorts; the bias is notably lower for the four cancers. The differential effect size (bias) of +0.05, corresponding to OR multiple equal 1.05 was reached for these LODs at control group age of 100 years; the bias age progression is displayed in Fig. S16 and Fig. S17 in Supplementary Chapters. The GWAS results are known to show underestimated SNP effect size for higher trait heritablities (Stringer et al., 2011; Banerjee et al., 2018), particularly relevant for AD and T2D, modeled here with *h*^2^ > 0.6—the heritability value when the underestimation effect becomes noticeable—with correspondingly large number of effect SNPs for common low-effect-size genetic architecture. Stringer et al. (2011) consider this phenomenon a facet of GWAS missing heritability characteristic to single SNP analysis. Multi-SNP analyses are proposed and are being developed (Zhu and Stephens, 2017; Banerjee et al., 2018). For the purposes of this study the customary single SNP analysis was sufficient for the relative bias determination for AD and T2D, which was found to closely follow the patterns of CAD and stroke.

The GWAS association simulations and age covariate adjustments and corresponding association standard errors are summarized in Table S8 in Supplementary Chapters. The progression shape in Fig. S17 in Supplementary Chapters implies that the bias is proportionate to a power function by age, and the bias magnitude progression appears proportionate to the effect size magnitude. The linear regression fitted the normalized effect bias according to Eq. (11) and Eq. (12). The quadratic bias adjustment, used by Chatterjee et al. (2003), resulted in reasonable bias adjustment, as seen in Fig. S18 in Supplementary Chapters. The best fit power of age regression, with the power exponent specific for each LOD, produced a better match, though the slight improvement over the quadratic regression will be likely indistinguishable in practical GWAS bias correction (compare Fig. S18 and Fig. S19 in Supplementary Chapters).

## Discussion

Performing a comprehensive set of validation simulations enabled the determination and generalization of the change in allele distribution with an increase in cumulative incidence for all genetic architectures described in the Methods section. The simulation results show that, for all genetic architectures, the change in the PRS depends on the cumulative incidence and the magnitude of heritability. When the same level of cumulative incidence is reached, the difference in allele distribution between diagnosed cases and the remaining unaffected population is identical. It therefore depends on the value of the cumulative incidence, and not on the incidence pattern that led to the achievement of a particular cumulative incidence value in the validation scenarios.

Next, the simulations were performed using the incidence rate patterns and model genetic architectures of each analyzed LOD, determining changes in the allele distribution with age and the resulting impact on GWAS discovery power. These results compared well with the findings from clinical, GWAS, and familial heritability studies, which are summarized below.

There are numerous reports of heritability, clinical predictive power, and GWAS discovery power diminishing with age for these LODs. Almgren et al. (2011) observed T2D heritability equal to 0.69 for the 35–60 year age-of-onset group, and negligible heritability for older ages. GWASs (de Miguel-Yanes et al., 2011; Mühlenbruch et al., 2013) segregating T2D risk SNPs by age have found that the risk factor values are higher for those under the age of 50, compared to the older cohorts. Regarding the variant types that are most likely associated with T2D, Fuchsberger et al. (2016) found that, with a high degree of certainty, they were able to attribute T2D liability to common variants rather than rare, high-effect variants. A cardiovascular disease (myocardial infarction) study by Nielsen et al. (2013) found the predictive power of parental history to decline for ages older than 50. Schulz et al. (2004) found familial history to be the best predictor of ischemic stroke for individuals under the age of 60. A review based on Framingham’s study (Seshadri et al., 2010) found the parental predictive power of stroke to diminish for those aged over 65. The heritability of Alzheimer’s disease has been estimated at 80% from twin studies (Naj and Schellenberg, 2017); Gatz et al. (2006) found heritability to be 79% at approximately 65 years of age, diminishing with increasing age. GWASs (Tan et al., 2013; Shen and Jia, 2016; Naj and Schellenberg, 2017) have come to similar conclusions. In summary, the predictive power of familial history for the above LODs is greatest for younger ages, specifically <65 years of age for AD, <50 for CAD, <60 for stroke, and <50 for T2D.

For the above LODs, the simulation results show high PRSs for the earliest-diagnosed cases. The risk allele case/control difference and the PRSs of newly diagnosed cases decrease rapidly with age progression. At a very old age, the individuals whose genotype would be considered low risk at an earlier age are the ones diagnosed with the disease; see Fig. 2. This also reinforces the validity of the clinical observation that the major risk factor for LODs is age itself.

The four cancers display a noticeably different pattern. The PRSs for the earliest-onset cases are lower than those for the above LODs, and this risk changes much less with age than for the above LODs. These results explain the observations of familial heritability studies: for three out of the four most prevalent cancers, twin studies have shown relatively constant heritability with age progression (Grönberg, 2003; Möller et al., 2016; Mucci et al., 2016; Graff et al., 2017). Determining the change in lung cancer heritability with age has proven somewhat more elusive (Hjelmborg et al., 2016), and no definitive conclusions have been published, largely due to the generally low documented heritability and substantial environmental component of this disease.

Prostate cancer is the only cancer that is somewhat controversial. Its heritability is reported at 57% by Hjelmborg et al. (2014), and prostate cancer reaches the highest maximum instance rate of the four most prevalent cancers reviewed. Therefore, according to the above observations and the results of the validation simulations, the relative MAF between cases and controls is likely to be higher than for other cancers. Nevertheless, the same article finds that the heritability of prostate cancer remains stable with age. It may be that this twin study result is somehow biased and that the heritability of prostate cancer is lower than stated in Hjelmborg et al. (2014), or perhaps this is a phenomenon specific to the populations or environmental effects of Nordic countries. Perhaps the earlier familial study (Grönberg, 2003), which estimated heritability at 42%, would be closer to the UK population incidence data used here. The verification simulation using a heritability of 42% produced results that matched more closely the patterns exhibited by the other cancers; see the resulting value shown in parentheses for the case multiple in Table 4. A more exhaustive literature investigation of the reviewed LODs is presented in Chapter S1 in Supplementary Chapters.

GWASs’ statistical discovery power is impaired by the change in individual distribution of the PRS at older ages. A larger number of cases and controls is needed at older ages to achieve the same statistical discovery power. The first four LODs, which exhibit higher heritability and cumulative incidence compared to cancers, require an increased number of participants in case/control studies for older ages. The cancers show a small increase in the number of participants required to achieve the same statistical power.

Individual values analysis, in which the individuals diagnosed each year are compared to all remaining healthy individuals, shows a rapid increase in the number of cases hypothetically needed to achieve the same statistical power, but this scenario would be impractical for a clinical study. The age-matched cohort studies benefit from the fact that the diagnosed individuals are accumulated from the youngest onset to the age of becoming a case in the cohort study, as well as being averaged over the cohort age range, resulting in a more moderate increase in the number of participants required, or a slower decline in GWAS discovery power for older cohorts. Age-matched cohort studies would require 1.5–2.1 times more participants at age 80 compared to the youngest possible age-matched cohorts in the case of stroke, CAD, AD, and T2D.

Designing cohorts composed of the youngest possible cases and the oldest available controls improves GWAS discovery power due to larger difference in risk allele frequency between cases and controls. This improvement leads to a lower number of participants being needed for GWASs when applied to the highest cumulative incidence and heritability LODs—so much so that about 50% fewer participants are required to achieve the same GWAS statistical power when control cohorts between 90 and 100 years of age are matched to the youngest case cohorts, with the reverse being the case with older age-matched cohorts. In this scenario, notably (20–25%) fewer participants are also needed to achieve the same statistical power in cancer GWASs, including those focusing on lung cancer.

Using non-age-matched cases and controls in GWAS cohorts, while improving the discovery power, may report higher than “true” association effect, which is expected with the enhanced difference in the effect SNP frequency between these cohorts, and would require appropriate adjustment, as demonstrated in the Results section. This study simulations imply that the adjustment may be simplified by the fact that the bias magnitude was found proportionate to the associated SNP effect size. Many GWAS association software packages offer automated covariate bias correction (Bhattacharjee et al., 2011; SAS Institute Inc, 2013; Conomos and Thornton, 2016; Harrell Jr et al., 2018; Purcell and Chang, 2019); the required bias adjustments may require addditional scripting in all packages.

Not every GWAS will be able to find a sufficient number of young cases, corresponding in this study to prevalence of approximately 0.25% for each LOD. However, due to a close to exponential rise in the incidence rate with age for most LODs, the case cohorts can be formed at a somewhat older age with correspondingly somewhat smaller improvement in GWAS discovery power. For all analyzed here LODs a majority of population would remain disease-free at ages 80- and 90-years old, with sufficient survivorship providing a large pool of older controls.

## Conclusions

This research was conducted with the goal of establishing whether any of the observational phenomena, including decreasing heritability with age for some notable LODs and the limited success of LOD GWAS discovery, can be explained by changes in the allele proportions between cases and controls due to the higher odds of more-susceptible individuals being diagnosed at an earlier age.

The simulation results reported above show that these phenomena can indeed be explained and predicted by the heritability of the LODs and their cumulative incidence progression. By simulating population age progression under the assumption of relative disease liability remaining proportionate to individual polygenic risk, it was confirmed that individuals with higher risk scores will become ill and be diagnosed proportionately earlier, bringing about a change in the distribution of risk alleles between new cases and the as-yet-unaffected population in every subsequent year of age. With advancing age, the mean polygenic risk of the aging population declines. The fraction of highest-risk individuals diminishes even faster. While the number of most-susceptible individuals and the mean population susceptibility both decline, the incidence of all LODs initially grows exponentially, doubling in incidence every 5 to 8.5 years (see the Methods section) and remains high at older ages, leading to a high cumulative incidence for some LODs. The increasing incidence rate in the face of declining polygenic risk for the as-yet-unaffected population can be explained as a consequence of the aging process, which itself is the major risk factor for LODs. In old age, people who have low genetic or familial susceptibility are increasingly becoming ill with an LOD.

LODs with low cumulative incidence and low familial heritability produce smaller changes in the allele distribution between affected individuals and the remaining population. The most-prevalent cancers are reported to have stable heritability with age, and therefore these GWASs are less affected by the increasing age of the participant cohorts. For these diseases, modification of the cohort age selection process by favoring younger cases and older controls will also lead to noticeable improvements in statistical power, albeit somewhat less prominent than for the higher-incidence and -heritability LODs.

Four of the most prevalent LODs—Alzheimer’s disease, coronary artery disease, cerebral stroke, and type 2 diabetes—exhibit both a high cumulative incidence at older age and high heritability. These simulation results show that a GWAS of any polygenic LOD that displays both high cumulative incidence at older age and high initial familial heritability will benefit from using the youngest possible participants as cases rather than age matching or statistically adjusting or compensating for age. In addition, cohort GWASs would benefit from using as controls participants who are as old as possible. This would allow for an additional increase in statistical discovery power due to the greater difference in risk allele frequency between cases and controls. For most LODs, ample numbers of still-unaffected individuals remain available at older ages to participate in the control cohorts.

## Supporting information

Supplementary Data: A zip file containing the simulation executable, the source code, R scripts, batch files, and simulation results.

## Acknowledgments

The author thanks Alexei J. Drummond at the University of Auckland for a number of helpful and challenging discussions.

## Supplementary material

**Supplementary Chapters** Supplementary chapters and figures are attached after the main manuscript text.

## Data availability

**Supplementary Data** A zip file containing the simulation executable, the source code, R scripts, batch files, and simulation results is provided on this site.

## SUPPLEMENTARY

### Supplementary Chapter S1: LOD heritability patterns with age based on familial and clinical studies and genome-wide association studies (GWAS)

The notion that the heritability of LODs always decreases with age is not entirely correct. A review of the clinical and familial studies and GWAS on the heritability of polygenic LODs within the typical age range of disease onset leads to a grouping of LODs into two broad categories: those with decreasing heritability with age and those with increasing or relatively constant heritability with age. Next, these categories are reviewed in detail, focusing primarily on the eight highly prevalent LODs analyzed in our simulations. These categories are used to organize the observational knowledge to enable the application of this knowledge to the main article’s simulations and the verification of the simulation results.

#### LODs with decreasing heritability with age

There is a large number of *highly environmentally affected LODs that exhibit decreasing heritability with age*. Three of these diseases carry some of the highest lifetime risk: coronary artery disease, cerebral stroke, and type 2 diabetes; see Table 1, summarized from Wienke et al. (2001), Zdravkovic et al. (2002), Devan et al. (2013) and Aparicio and Seshadri (2017).

**Supplementary Table 1.**
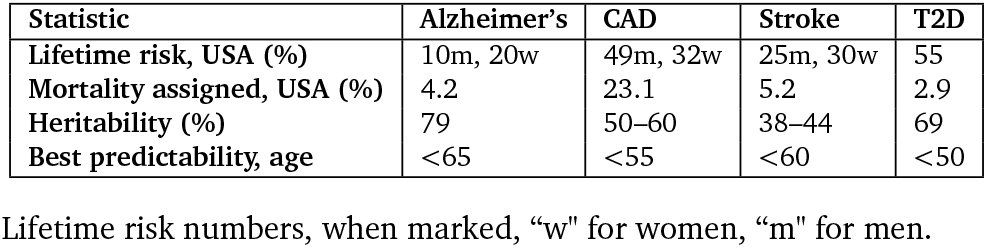
Population statistics of LODs characterized by decreasing heritability with age.

Falconer (1967) noted that *“the increase of incidence associated with a variable age of onset can be due to either an increase of the mean liability or an increase of the variance of liability. Consideration of the changes of liability that individuals may undergo as they grow older shows that an increase of variance with increasing age is to be expected, and since the additional variance is likely to be mainly environmental, a reduction of the heritability is to be expected*.” Falconer further pointed out that *“the heritability of liability to diabetes, estimated from the sib correlation, decreases with increasing age. For people under 10, heritability is about 70 or 80%, and it drops to about 30 or 40% in people aged 50 and over. The decrease of the heritability is attributable to an increase of environmentally caused variation. The increased environmental variation is not enough to account in full for the increasing incidence, so there is probably also an increase of the mean liability with increasing age*.”

In the 1960s, the distinction between autoimmune Mendelian type 1 diabetes and late-onset polygenic type 2 diabetes (T2D) was not known, but it was suspected that there may be two distinct mechanisms. However, this conclusion—of an increase in liability with age, and accordingly blurred heritability—is observed for T2D as well as other LODs.

The greatest heritability for T2D is observed in the 35–60 (0.69) year age of onset group, (Almgren et al., 2011) and heritability declines to only 0.31 when the upper age limit is increased to 75 (making the age range 35–75). In the over-60 group, the “environmental” component is the primary cause of new T2D cases. The environmental component in this case includes systemic and tissue-specific deterioration with age and the cumulative external environmental effects with increased time duration. Just as Falconer did 60 years earlier, this study concludes that T2D heritability decreases with age and that liability may be more accurately predicted in younger individuals. One review (Talmud et al., 2014) cites two studies that corroborate this view. The first concluded that recalculating the genetic risk for T2D by splitting a cohort by age below and above 50 years using 40 T2D risk SNPs finds that the risk factor values are higher in the younger group (de Miguel-Yanes et al., 2011). Meanwhile, Almgren et al. (2011) correlated the heritability and familiality of T2D with quantitative traits and found a very significant drop in heritability over the age of 60.

The conclusion is that, for reliable GWASs, younger is better: T2D patients under the age of 60—or, even better, under the age of 50—should be chosen. Regarding the variant types that are most likely associated with T2D, Fuchsberger et al. (2016) found that they were able, with a high degree of certainty, to attribute T2D liability to common variants rather than rare, high-effect variants.

Nielsen et al. (2013) cardiovascular disease (myocardial infarction) study provides implicit confirmation of decreasing heritability with age. The predictive power of parental history is as follows: paternal relative risk (RR) = 3.30 for ages <50 and 1.83 for ages >50; maternal RR = 3.23 for ages <50 and 2.31 for ages >50.

Schulz et al. (2004) found familial history to be the best predictor of **ischemic stroke** for individuals under the age of 60, with an overall odds ratio (OR) of 1.73. Relative OR, compared to the under-60 cohort, was 0.95 for the 60–70 age band and 0.77 for individuals over the age of 75.

A review based on Framingham’s study (Seshadri et al., 2010) supplies very useful information about parental history of stroke. Even though the grouping used on the parental side is stroke under 65, on the descendant side, there are statistics showing RR both below and above the age of 65. For descendants whose parents had a stroke before the age of 65, the stroke RR was determined. Overall, the RR was 3.79 under the age of 65 and 2.21 over the age of 65; the HR for ischemic stroke was 5.45 under the age of 65 and 2.47 over the age of 65. Additional implicit information from this data, which supports the same conclusion, is listed in Allport et al. (2016)

The heritability patterns for these diseases are summarized in Table 3. There is qualitative and, increasingly, quantitative knowledge about the progressively declining heritability of these diseases at ages above 50, as well as the decreasing associated familial and GWAS predictive power; see Nielsen et al. (2013), Schulz et al. (2004), Seshadri et al. (2010), Bevan et al. (2012), Devan et al. (2013) and Fuchsberger et al. (2016) These studies found familial history to be the better predictor of next-generation disease only when the participants in the parental generation are relatively young; see de Miguel-Yanes et al. (2011), Talmud et al. (2014), Almgren et al. (2011) and Table 1.

An environmental effect on the heritability of cardiovascular disease and T2D with age is evident, (Falconer, 1967; Poulsen et al., 1999) including influences such as spousal environment (Jee et al., 2002).

In addition, T2D is a major co-morbidity factor for CAD and cerebral stroke, as well as causally correlated adiposity and hypertension, which are by themselves associated with CAD and cerebral stroke and other LODs. In the presence of T2D, these diseases develop years and even decades earlier than the typical onset ages (Boehme et al., 2015). For instance, twin studies on the heritability of BMI (a co-morbidity often preceding T2D) show the highest heritability of 85% at 18 years of age, after which heritability slowly declines throughout the lifespan (Elks et al., 2012).

It must be noted that the majority of diseases are influenced to various degrees by environmental factors. The three diseases just reviewed show incomparably higher environmental influence than Alzheimer’s disease (AD). For AD, neither lifestyle nor painstakingly developed medications can markedly influence the progression of the disease. In contrast, CAD, cerebral stroke and T2D are often considered by the medical community to be primarily influenced by lifestyle and environment (Lloyd-Jones et al., 2006; Mahmood et al., 2014; Boehme et al., 2015; Diapedia: Epidemiology of type 2 diabetes).

In conclusion, the highly prevalent LODs exhibiting high environmental correlation with onset ages also show decreasing heritability with age. This is combined with an exponential increase in incidence with age. In the case of CAD and cerebral stroke, the exponential incidence rate increase proceeds beyond 80 years of age.

Another type of *LOD showing heritability that declines with age can be described as a mode of failure with aging.* Alzheimer’s disease begins relatively late, but from there, its incidence rises exponentially to extremely old age (Brookmeyer et al., 1998). The heritability of Alzheimer’s disease is estimated at 80% from twin studies (Naj and Schellenberg, 2017); both familial studies and GWAS estimate heritability at 79% Gatz et al. (2006) at approximately 65 years of age, diminishing with increasing age. Tan et al. (2013); Shen and Jia (2016); Naj and Schellenberg (2017)

A clinical study documenting the association between the APOE genotype and Alzheimer’s disease (Farrer et al., 1997; Davidson et al., 2007) reports the change in odds ratio with age of APOE e4/e4 and APOE e3/e4 carriers, which is summarized for the Caucasian population in Table 2.

**Supplementary Table 2.** Alzheimer’s disease odds ratio by age and APOE alleles, relative to e3/e3 allele carriers. Values summarized from Farrer et al. (1997).

| APOE allele / Age (y) | 55 | 60 | 65 | 70 | 75 | 80 | 85 | 90 |
| --- | --- | --- | --- | --- | --- | --- | --- | --- |
| e4/e4 OR | 14.1 | 15.0 | 14.3 | 12.1 | 9.5 | 6.1 | 3.7 | 2.0 |
| e4/e3 OR | 3.5 | 3.7 | 3.8 | 3.6 | 3.3 | 2.7 | 2.3 | 1.7 |

Another review (Naj and Schellenberg, 2017) concludes that the typical age at onset is 68.8 years for APOE e4/e4 carriers, 75.5 years for e3/e4 carriers, and 84.3 years for carriers without e4. Moreover, the APOE e4 effect is age dependent, giving a broad-stroke assessment that the e4 allele effect is most prominent between the ages of 60 and 79 and gradually diminishes after the age of 80. This fits well with the assessment (Farrer et al., 1997) summarized in Table 2.

Table 3 summarizes the information in the literature about the decreasing heritability of the LODs referenced above.

The model presented by Brookmeyer et al. (1998) hypothesized that, if the AD incidence curve could be delayed by five years, the overall prevalence of AD would be half the projected rate, assuming unchanged mortality from other causes. AD prevalence in this study is limited by applying a 1.4 mortality multiplier to AD patients compared with the unaffected population.

**Supplementary Table 3.** Heritability and risk statistics for LODs exhibiting decreasing heritability with age.

| Disease | Heritability/risk, younger age | Heritability/risk, older age |
| --- | --- | --- |
| AD e3/e4 (Farrer et al., 1997) | OR=3.8, 65y | OR=1.7, 90y |
| AD e4/e4 (Farrer et al., 1997) | OR=15.0, 60y | OR=2.0, 90y |
| CAD paternal (Nielsen et al., 2013) | RR=3.30, < 50y | RR=1.83, > 50y |
| CAD maternal (Nielsen et al., 2013) | RR=3.23, < 50y | RR=2.31, > 50y |
| Stroke (Schulz et al., 2004) | OR=1.63, < 60y | OR=0.77, > 70y |
| Stroke all (Seshadri et al., 2010) | RR=3.79, < 65y | RR=2.21, > 65y |
| Stroke ischemic (Seshadri et al., 2010) | RR=5.45, < 65y | RR=2.47, > 65y |
| T2D (Almgren et al., 2011) | $h^2 = 0.69$, 35-60y | $h^2 = 0.31$, 35-75y |

OR = odds ratio; RR = relative risk; $h^2$ = heritability

While AD progression is difficult to influence with lifestyle changes or medications, AD incidence at comparable ages has decreased by about 30% since the 1980s in many Western countries (Binder and Schumacher, 2016; Wu et al., 2017) due to undetermined causes. As life expectancy increases, AD lifetime incidence and prevalence are expected to regain ground.

In conclusion, AD shows an exponentially increasing incidence rate up to the most advanced ages, while also displaying heritability that declines with age.

#### LODs exhibiting stable heritability with age

LODs with relatively constant heritability with age and infrequent types of LODs with increasing heritability with age are grouped in this category. As found in the reviewed literature, the increase in heritability, when observed, is moderate. The diseases showing *slightly increasing heritability with age* are found to be those affecting the skeletal system, for instance, osteoarthritis, particularly of large joints such as the hip or lower back. One study (Skousgaard et al., 2015) shows that both the incidence and heritability of advanced osteoarthritis of the hip and lower back increase with age.

It is evident that younger cases are more environmentally and less genetically correlated. For example, osteoarthritis at a younger age is often due to trauma rather than genetics (Amoako and Pujalte, 2014; Warner and Valdes, 2016). At the age of 60, the influence of genetic and environmental components is roughly equivalent, and by the age of 70, heritability increases to 75% and stays close to this level into the 90s. Heritability is even higher and increases with advanced age for osteoarthritis of the spine at multiple locations (Spector and MacGregor, 2004).

The increase in heritability for these diseases is seen to be relatively modest and extends from an initially high level. Many osteoarthritis-affected structures and corresponding diagnoses, with different ages of maximum incidence and heritability by sex and age, do not follow this pattern (Skousgaard et al., 2016).

The osteoporosis findings are similarly varied, with studies finding no heritability of pathology for some bone structures and strong heritability for others (Ralston and Uitterlinden, 2010). Specifically, the osteoporosis associated with bone breaks is very heritable and shows a slight increase in heritability into older age (Shaffer et al., 2008). This is explicable by the fact that, for osteoporosis, the main risk component—the shape and size of the bone—is strongly heritable. Genetics in this case determines the early developmental stages of an organism, when the structures take shape. Similar reasoning applies to osteoarthritis, which is related to defects in collagen and connective tissue formation. The malignancy occurs after many decades of life, when wear, deterioration and diminishing repair capacity cross the threshold leading to pathology.

In conclusion, some LODs with their roots in the early development of an organism’s structures may display strong heritability late in life and even increasing diagnostic heritability as aging progresses. GWAS has found only a small set of SNPs that provides very limited risk prediction for these diseases (Loughlin, 2015; Warner and Valdes, 2016). Apparently, the research cannot be impeded by the increasing heritability with age of the GWAS cohorts. *Relatively stable heritability with advancing age is a distinguishing feature of cancers*. Accurate information about heritability at different ages is not sufficiently explored for most cancers. Fortunately, during this decade, a number of studies have shed light on the age-related heritability of three out of the four most prevalent cancers, and these data allow us to extrapolate the expectations to the fourth: lung cancer.

The lifetime risk of developing any type of cancer in the US is 38% for women and 40% for men, (Lifetime Risk of Developing or Dying From Cancer) and the 2016 fraction of mortality directly attributed to cancer was 21.8%, the second-highest after heart disease (Murphy et al., 2017). In the UK, the corresponding numbers are higher, at 47% and 53%, respectively, (Ahmad et al., 2015; Cancer Statistics for the UK) with the higher likelihood perhaps attributable to the UK’s longer life expectancy. Each specific type of cancer constitutes a small fraction of overall lifetime risk, with breast, prostate, lung, and colorectal cancer being the four most prevalent.

Next, the latest heritability and incidence research for these four cancers is summarized.

##### Breast cancer (BC)

Breast cancer (BC) is well researched, with studies delving into all aspects of BC. Like prostate cancer, the two largest genetic predictors of BC are mutations in the BRCA1 and BRCA2 genes. The BRCA1/2 genes are involved in the homologous repair of double-stranded DNA breaks, working in combination with at least 13 known tumor suppressor proteins (Haley, 2016). Defects in the BRCA1/2 proteins disable homologous double-stranded DNA break repair, and the cell falls back on the use of imprecise non-homologous repair mechanisms; this leads to the accumulation of mutations, eventually leading to cancer. BRCA1/2 mutations are the most important predictor of breast cancer. The review by Haley (2016) states that the frequency of BRCA mutations varies with geographic location and ethnicity, ranging from a 0.02% mutation carrier rate in some populations to 2.6% in the Ashkenazi Jewish population due to ancient founder mutations. Other founder mutations have been reported in the Dutch, Swedish, French Canadian, Icelandic, German, and Spanish populations. In On-tario, Canada, for instance, the frequency of mutation carriers is 0.32% for BRCA1 and 0.69% for BRCA2 (Risch et al., 2006).

An early study (Ford et al., 1998) analyzing families with at least four cases of BC found that the disease was linked to BRCA1 in 52% of cases and BRCA2 in 32% of cases (with only 16% remaining for other causes). Taking into account ovarian cancer in addition to BC resulted in 81% of cases being due to BRCA1, while 76% of cases in families with both male and female BC were due to BRCA2. The lifetime risk of BC for women both in the US and the UK is 12% (Lifetime Risk of Developing or Dying From Cancer; Cancer Statistics for the UK). As Haley (2016) summarized, carriers of BRCA1 have a lifetime risk of developing BC equal to 60–70%, and an additional 40% risk of developing ovarian, fallopian, or primary peritoneal cancers. For BRCA2 carriers, the risks are 45–55% for BC and 25% for ovarian cancer. These numbers closely correspond to the aforementioned study (Ford et al., 1998). Möller et al. (2016) presented in-depth data on the heritability by age of breast and ovarian cancer for BRCA1/2 carriers. The study demonstrated that the genetic liability, while exhibiting a slight downward trend, remains relatively constant and exceeds the common environmental component at all ages.

One of the most recent studies (Kuchenbaecker et al., 2017) provides further clarification, stating that BC incidences increase rapidly in early adulthood until the ages of 30 to 40 for BRCA1 carriers and until the ages of 40 to 50 for BRCA2 carriers, thereafter remaining at a relatively constant incidence rate of 2–3% per year until at least 80 years of age; see Table 4. This study’s calculations based on this data show that the initial increase in incidence is exponential before flattening into the constant horizontal incidence rate approximation; a logistic approximation also fits. The exponential doubling rate, until it reaches the constant incidence level, is also consistent with all other diseases reviewed, showing an incidence doubling time of five years for BRCA1 and eight years for BRCA2 (the BRCA1 calculation, based only on two data points, is less accurate). A much earlier review study (Antoniou et al., 2003) collected the same kind of statistics as Kuchenbaecker et al. (2017) and arrived at similar conclusions.

**Supplementary Table 4.** BRCA1/2 carriers incidence rate by age, data from Kuchenbaecker et al. (2017)

| Gene | ≤20 | 21–30 | 31–40 | 41–50 | 51–60 | 61–70 | 71–80 |
| --- | --- | --- | --- | --- | --- | --- | --- |
| BRCA1 (%) | 0 | 0.59 | 2.35 | 2.83 | 2.57 | 2.50 | 1.65 |
| BRCA2 (%) | 0 | 0.48 | 1.08 | 2.75 | 3.06 | 2.29 | 2.19 |
| BRCA1 cum. risk (%) | 0 | 4 | 24 | 43 | 56 | 66 | 72 |
| BRCA2 cum. risk (%) | 0 | 4 | 13 | 35 | 53 | 61 | 69 |

Möller et al. (2016) study found a somewhat lower life-time BC risk of 8.1% in Nordic countries compared to 12% in the US and estimated heritability at 31%.

In addition to BRCA1/2, Mavaddat et al. (2010) and Haley (2016) also list a number of high-penetrance gene mutations—the TP53, PTEN, STK11, and CDH-1 gene mutations—giving a lifetime probability of cancers in general of about 90% and specifically a female breast cancer probability above 50%.

Several rare gene mutations—CHEK2, PALB2, ATM, BRIP1CHEK2, PALB2, ATM, and BRIP1—are also associated with a breast cancer relative risk in the range of 1.5–5.0. In aggregate, these high-effect mutations are correlated with only approximately 10% of hereditary breast cancers (Risch et al., 2006; Haley, 2016).

To date, GWAS attempts to discover common polygenic variants of low effect size have had only limited success. One review study (Lyra-Junior et al., 2017) outlines the history and accomplishments of breast cancer GWAS over a decade of research. The most recent high-powered consortium study (Michailidou et al., 2017) included 122,977 cases and 105,974 controls of European ancestry as well as 14,068 cases and 13,104 controls of East Asian ancestry. The study verified 102 previously reported SNPs, finding that 49 of them were reproducible. The study also found that the majority of discovered SNPs reside in noncoding areas of the genome. The discovered set of polygenic SNPs allows for the explanation of approximately 4% of heritability on top of the 14% explained by known high-penetrance SNPs, bringing the predictive power to 18%. This GWAS estimated the familial heritability of breast cancer at 41%—a possible exaggeration, because it significantly exceeds the 31% estimated by Möller et al. (2016) and the 27% estimated by Mucci et al. (2016)

###### Breast cancer conclusions

The familial heritability studies and BRCA1/2 clinical studies show that breast cancer heritability is relatively constant over the age of 40 for both mutations. A number of high-penetrance gene mutations can explain an additional fraction of heritability, totaling 10–14%.

The GWAS described above (Michailidou et al., 2017) also found multiple SNPs located in non-coding areas to be correlated with the candidate gene promoters and activity modifier areas. This improves the possibility that the common variant component may be able to explain a larger fraction of heritability. It appears at this time, based on Möller et al. (2016) statistics, that breast cancer heritability for the polygenic component may also be relatively constant after the age of 40 or may only slightly decline with age.

##### Prostate cancer (PC)

The effects and risks of the BRCA1/2 genes and their mutations described in the breast cancer section apply in a very similar way to the incidence of PC.

A study by Lecarpentier et al. (2017) found that lifetime PC risks are approximately 20% for BRCA1 mutations carriers and 40% for BRCA2 mutation carriers, while, overall, BRCA1/2 is associated with only 2% of all PC cases. In addition, BRCA1/2 accounts for 10% of male breast cancer cases. The lifetime risk of male breast cancer in mutation carriers is estimated at 5–10% for BRCA1 mutations and 1–5% for BRCA2 mutation carriers. Therefore, compared to breast cancer, BRCA1/2 mutations are associated with a smaller fraction of heritability.

The lifetime risk of PC in men is estimated at 6% for Danish cohorts and 12% for Finnish, Norwegian, and Swedish cohorts. The lifetime risk of developing PC in the US and the UK is 12% (Lifetime Risk of Developing or Dying From Cancer; Cancer Statistics for the UK). PC heritability has been estimated at 57% (Hjelmborg et al., 2014; Mucci et al., 2016) and 42% by an older study (Grönberg, 2003). The Nordic twin study (Hjelmborg et al., 2014) presents strong evidence that the heritability of PC remains stable or even slightly increases between the ages of 65 and 100. As with breast cancer, the fraction of PC attributed to highly malignant mutations is low. Known rare, high-effect-size variants such as BRCA1/2, ATM, and HOXB13 explain only 10–12% of heritability (Wu and Gu, 2016; Mancuso et al., 2016; Walsh, 2017; Lecarpentier et al., 2017). Recently, Eeles et al. (2017) using an imputed meta-analysis for 145,000 men, reported that the GWAS polygenic score they obtained explains 33% of the familial relative risk.

Wu and Gu (2016) concluded that the search for the missing heritability may be better served by high-coverage whole-genome sequencing (WGS); however, due to the cost and complexity, it is not currently feasible to obtain this much high-quality data. In the absence of more predictive genetic data, Wu and Gu (2016) noted that the best predictor of PC is age itself.

###### Prostate cancer conclusions

The conclusions for PC heritability are very much the same as for breast cancer. While the heritability is higher than that of BC, it appears even more likely to remain constant or slightly increase with age, notwithstanding the smaller number of known rare, large-effect-size mutations that can be used to explain the heritability of PC.

##### Colorectal cancer (CRC)

The lifetime risk of developing CRC in the US is 4.1% for women and 4.5% for men (Lifetime Risk of Developing or Dying From Cancer). In the UK, the corresponding numbers are 5% and 7% (Cancer Statistics for the UK).

The Nordic twin studies (Mucci et al., 2016; Graff et al., 2017) estimated CRC heritability at 40%. A number of studies have included separate classifications for colon cancer, with a heritability of 15%, and rectal cancer, with a heritability of 14%, while the combined percentage is more than double the individual ones. This example may indicate that, while subdivisions exist in the medical diagnoses that may make a difference for surgical or treatment purposes, and while even the carcinogenicity manifestations may differ between subareas of the organ, from the perspective of the heritability of the liability, they are inherited as a single condition.

CRC heritability is also relatively constant between the ages of 50 and 95 in twin studies (Graff et al., 2017). Compared to the two previously reviewed cancers, there is a larger number of identified predisposing mutations and syndromes, such as Lynch syndrome, familial adenomatous polyposis, Peutz–Jeghers syndrome, juvenile polyposis syndrome, MUTYH-associated polyposis, NTHL1-associated polyposis, and polymerase proofreading-associated polyposis syndrome (de Voer et al., 2016; Jiao et al., 2014).

Graff et al. (2017) study concluded that, although a small number of genetic variants have a substantial effect on CRC, a considerable portion of its heritability is thought to result from multiple low-risk variants. de Voer et al. (2016)) concurred that penetrant high-effect gene variants are found in 5–10% of CRC cases. A GWAS review (Schmit et al., 2016) found that more than 50 SNPs have been identified as credibly associated with CRC risk, yet these only account for a small proportion of heritability. In GWAS, common, genome-wide variants are able to account for 8% of heritability.

###### Colorectal cancer conclusions

The conclusions are much the same as for BC and PC.

##### Lung cancer (LC)

The lifetime risk of developing LC in the US is 6.0% for women and 6.9% for men (Lifetime Risk of Developing or Dying From Cancer). In the UK, the corresponding numbers are 5.9% and 7.6% (Cancer Statistics for the UK).

The LC pattern of heritability is not easy to ascertain. According to Kanwal et al. (2017) approximately 8% of lung cancers are inherited or occur as a result of a genetic predisposition. The Nordic twin studies review (Mucci et al., 2016) estimated the heritability of LC at 18% (within a likely range of 0–42%). Heritability studies require controlling for environmental factors, particularly tobacco smoking. It is perhaps for this reason that the Nordic twin studies consortium, which was invaluable in the three other cancer analyses, primarily restricted itself to analyzing the effects of tobacco smoking on LC (Hjelmborg et al., 2016).

Factors such as asbestos, industrial smoke and pollutants, high levels of domestic radon in some areas of the world, or exposure of miners to radon or other sources of radiation may influence incidence and, if not accounted for, may affect heritability estimates (Krewski et al., 2005; Carr et al., 2015; Malhotra et al., 2016). Hereditary mutations of genes that regulate DNA repair, including BRCA1/2, TP53 and others, also increase the risk of LC, as with almost any cancer (Kanwal et al., 2017).

Due to the low heritability of LC, GWASs’ success at identifying predictive common SNPs has been limited (Weiss-feld et al., 2015). Some studies explain part of the LC incidence by reference to causal epigenetic effects (Shi et al., 2017). The heritability value of 18% given by Mucci et al. (2016) has a very broad range. An earlier study (Yang et al., 2013) noted that tobacco smoking is by far the largest causal factor for LC, and the heritability of smoking itself may outweigh any other LC heritability.

Mucci et al. (2016) also considered smoking, but the high value reported by them exceeds the previous consensus and may need further corroboration. LC perhaps belongs to the difficult-to-analyze, non-additive traits of heritability noted by Polderman et al. (2015). This study considers LC heritability to be closer to 10%.

###### Lung cancer conclusions

In conclusion, an age-related heritability pattern for LC is lacking, and while it is impossible to make definitive conclusions, it can be hypothesized that LC follows a similar pattern to the other three cancers reviewed.

In summary, the heritability patterns of cancers were not systematically investigated until relatively recently. A small number of familial studies (Hjelmborg et al., 2014; Möller et al., 2016; Haley, 2016; Graff et al., 2017) and a more recent study that is particularly informative about the incidence of BRCA1/2 mutations with age (Kuchenbaecker et al., 2017) have finally allowed researchers to determine that cancer heritability remains relatively constant with age. Table 5 summarizes the findings from the reviewed literature in relation to breast, prostate, colorectal, and lung cancer. Studies ascertaining the heritability of lung cancer with age are absent from the literature; data may be difficult to collect due to the relatively low heritability of the disease.

Most lung cancer incidence is environmental, and lung cancer does not have specific, highly malignant mutations that may cause a noticeable fraction of heritability. The mostly polygenic fraction of lung cancer heritability is hypothesized to be similarly stable with age, as is the case with the other three cancers reviewed.

**Supplementary Table 5.** Patterns of heritability by age for most common cancers.

| Cancer type: | Breast | Prostate | Colorectal | Lung |
| --- | --- | --- | --- | --- |
| Lifetime risk, USA (%) | 12 | 12 | 4.5m 4.1w | 6.9m 6w |
| Heritability (%) | 31 | 57 | 40 | 8–18 |
| Incidence from highly detrimental mutations (%) | 10–14 | 10–12 | 5–10 | minor |
| Polymorphic incidence (%) | 86–90 | 88–90 | 90–95 | major |
| Heritability trend (50y–100y) | flat / slight decline<br>(Ford et al., 1998; Antoniou et al., 2003; Risch et al., 2006; Mavaddat et al., 2010; Mucci et al., 2016; Haley, 2016; Möller et al., 2016; Kuchenbaecker et al., 2017; Michailidou et al., 2017) | flat / slight incline<br>(Hjelmberg et al., 2014; Wu and Gu, 2016; Mancuso et al., 2016; Walsh, 2017; Lecarpentier et al., 2017; Eeles et al., 2017) | flat<br>(Jiao et al., 2014; Schmit et al., 2016; de Voer et al., 2016; Graff et al., 2017) | likely flat<br>(Krewski et al., 2005; Weissfeld et al., 2015; Carr et al., 2015; Malhotra et al., 2016; Mucci et al., 2016; Hjelmberg et al., 2016; Kanwal et al., 2017; Shi et al., 2017; Wang and Wang, 2017) |

Lifetime risk numbers, when marked, "w" for women, "m" for men. Lifetime risk numbers, when marked, "w" for women, "m" for men.

Because cancer development is primarily a consequence of mutations and epigenetic effects leading to unconstrained propagation of the clonal cell population, in the long term, cancers are inevitable for most multicellular organisms, including humans (Marusyk and DeGregori, 2008; Tomasetti and Vogelstein, 2015; Guedj et al., 2016; Ribezzo et al., 2016; Nelson and Masel, 2017).

Due to cancer’s constant heritability with age, the effect of age is likely to be insignificant for GWASs’ discovery of cancer polygenic scores and their corresponding predictive power. This could also apply to any LOD that follows a similar heritability pattern, that is, one that is relatively constant with age.

### Supplementary Chapter S2: Incidence functional approximation used in preliminary validations

To determine the effect of disease incidence with age progression on allele frequencies in the population and the difference in allele frequency between the newly affected and remaining unaffected populations, three incidence dependencies with age were used.

1. Constant incidence:

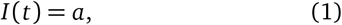

where *a* is a constant representing a horizontal line. Yearly incidence values of 0.0015, 0.005, and 0.02 (0.15% to 2%) were selected.
2. Linear incidence:

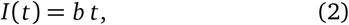

where *b* is a slope of the linear progression with intercept 0. Slope values of 0.003, 0.01, and 0.04 were selected. This means that incidence begins at 0 and increases to an incidence equal to 0.3%, 1%, and 4%, respectively, at 100 years of age to match the cumulative incidence of 1) above. These values were chosen to simplify the evaluation via simulation. The simulation was run with zero mortality, and the values were chosen to keep cumulative incidence at the same level—0.44 (44%)—at 100 years of age for the highest of either the constant or linear incidence progression.
3. In addition, an evaluation exponential incidence progression was used:

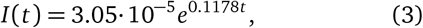

fitted to achieve a similar cumulative incidence at the most advanced age.

In all five scenarios described in the main article, the values of the case and control means and standard deviation/variance are identical when the cumulative incidence reaches the same level.

Two heritability scenarios were validated: 30.5% and 80.5%; see Table 6.

**Supplementary Table 6.** Linear and constant incidence validation scenarios.

|  | A | B | C | D | E |
| --- | --- | --- | --- | --- | --- |
| Scenario 1. Variants: | 400 | 625 | 1375 | 50 | 25 |
| Achieved heritability: | 0.3068 | 0.308 | 0.3075 | 0.296 | 0.3142 |
| Scenario 2. Variants: | 3725 | 5850 | 12775 | 500 | 225 |
| Achieved heritability: | 0.8047 | 0.8064 | 0.8049 | 0.8078 | 0.8048 |

The target heritability is 0.305 (30.5%) for validation scenario 1 and 0.805 (80.5%) for validation scenario 2 due to the genetic architecture model requiring multiples of 25 variants.

#### Validating allele distribution change in model genetic architectures using systematic incidence progressions

A set of validation simulations was run to verify the behavior of the model genetic distributions for the three types of incidence progression described above. The validation simulations based on the constant, linear and exponential incidence rates confirmed that both of the mean polygenic scores, for the population and for the cases, viewed in the individual values analysis for each age depend on the cumulative incidence and the magnitude of heritability, with neither being dependent on the shape of incidence progression with age.

From the validation simulations, the cumulative incidence, regardless of the incidence progression pattern, was found to produce a virtually identical polygenic score distribution for cases and the remaining unaffected population; see the genetic common allele low effect size plotted in Supplementary Fig. 2.

Between the genetic architectures, there is also a relatively small difference in the polygenic scores of the population and the cases; see Supplementary Fig. 3. As can be seen, the low-effect-size scenarios A, B, and C, progressing in allele frequency from common to rare, are practically indistinguishable from each other.

The higher-effect-size architectures (D and E) show a slightly larger fraction of higher-polygenic-score individuals or, more precisely, a slightly larger representation of higher- and low-polygenic-score individuals. The qualitative picture is close to identical among all five scenarios.

### Supplementary Chapter S3: LOD incidence functional approximation

The simulations were applied to eight of the most prevalent LODs: Alzheimer’s disease, type 2 diabetes, coronary artery disease, and cerebral stroke, and four late-onset cancers: breast, prostate, colorectal, and lung cancer. First, the functional approximation of the clinical incidence data used for the simulations is described. The incidence progression of the LODs with age is presented in Supplementary Fig. 1. The initial incidence rate (the fraction of the population newly diagnosed each year) increases exponentially with age. This exponential growth continues for decades, after which the growth in older cohorts may flatten, as in the case of T2D (Boehme et al., 2015). In the case of cerebral stroke and CAD, the clinical studies indicate a slowdown of the incidence for individuals over the age of 85; (Rothwell et al., 2005) accordingly, a constant level was used for the exponential approximation Eq. 4.

The incidence of Alzheimer’s disease, on the other hand, continues exponentially past the age of 95, reaching incidences above 20% (Brookmeyer et al., 1998). Cancer progression reaches only a small fraction of the incidence levels of the above-mentioned LODs, even for the four most prevalent cancers. Generalizing to other cancers, the incidence is much lower for more than a hundred of the less prevalent cancer types.

**Supplementary Fig. 1.**
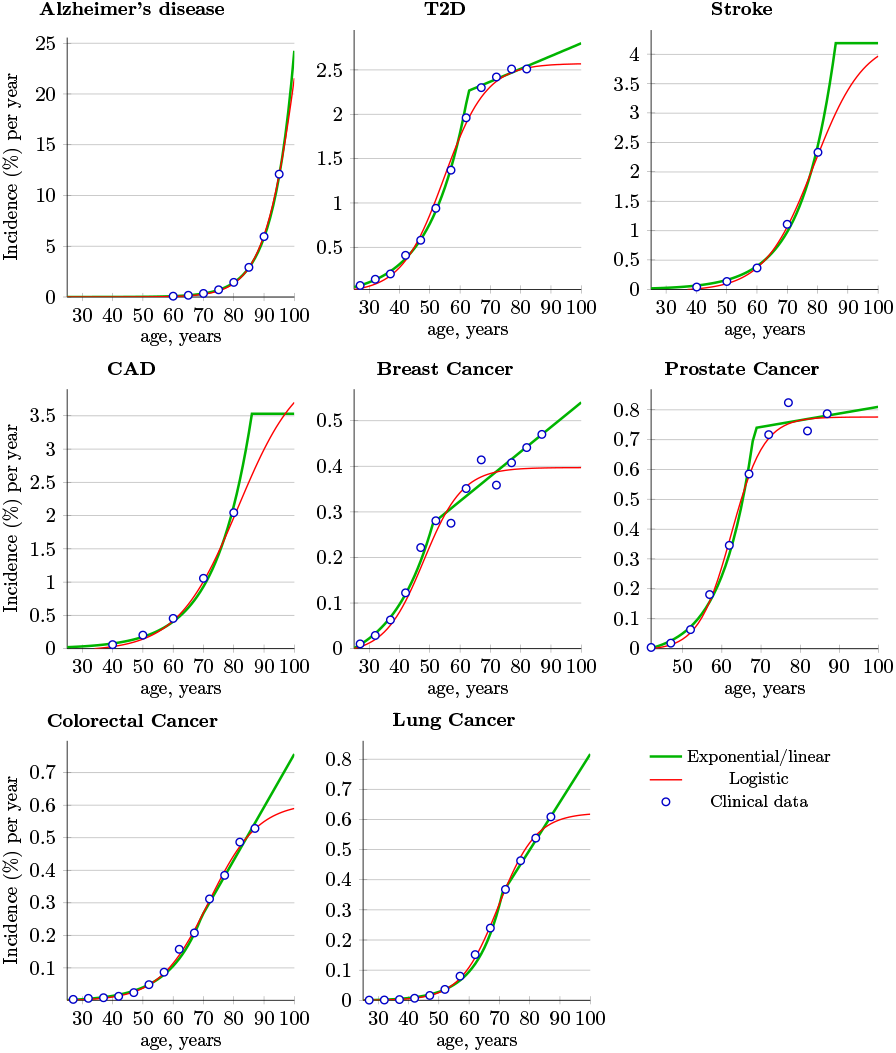
LOD clinical incidence rates and functional approximations. Two functional approximations of clinical data: exponential followed by linear and logistic. The R script automating NLM (nonlinear model) regression for both approximation curves is available in Supporting Information.

To evaluate each LOD’s allele redistribution with age, it was necessary to approximate the yearly incidence from much rougher-grained statistics. An R script automated the determination of the best fit for logistic and exponential regression from the clinical incidence data. The script also calculated lifetime incidence from our functional approximations; it closely matched the disease clinical statistics presented in Tables 1 and 5.

The incidence approximation *I* (*t*) is represented mathematically by Eq. 4; *a*, *b*, and *c* are exponential approximation parameters, *i* and *s* are the linear regression intercept and slope, respectively, and *t* is time in years.

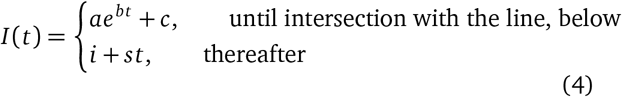

A logistic approximation of the clinical data is shown in red in Supplementary Fig. 1. It is characterized by the following equation:

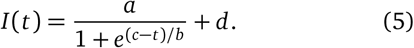

The incidence rate in the logistic curve slows faster than the incidence rate in the exponential curve and also approximates the incidence rate with age. It follows a similar pattern, with an initial exponential rise and a logistic inflection point occurring at quite advanced ages. Thus, the clinical data and corresponding approximations show the higher representation of older people in the patient cohorts.

For all LODs, decades-long initial exponential sections were observed in the incidence curve. The exponent conforms to a relatively narrow range of doubling the incidence rate, fitting between 5 and 8.5 years. While the absolute incidence rate differs significantly, the exponent constant multiplier *a*, which is equivalent to the linear regression intercept for *log*(*a*) in the *I* (*t*) function, mainly controls the rise, or the initial incidence onset, of the incidence rate; see Supplementary Fig. 1.

From this are found the logistic recursion inflection points at values shown in Table 7. The exponential incidence rise follows with high precision up to the ages shown in the table, and the rapid rise in the incidence rate continues past these ages.

**Supplementary Table 7.** Age to which LOD incidence rate rises exponentially.

|  | Highly prevalent LODs |  |  |  | Cancers |  |  |  |
| --- | --- | --- | --- | --- | --- | --- | --- | --- |
|  | AD | T2D | CAD | Stroke | Prostate | Colorectal | Breast | Lung |
| Age (years) | 103 | 55 | 81 | 79 | 48 | 62 | 72 | 70 |

The logistic approximation produced a good, simple fit for seven of the eight diseases. While the logistic approximation could also have been used for breast cancer, the exponential-plus-linear approximation showed a better fit and was therefore preferred.

As this paper makes extensive reference to the incidence of LODs, some of the commonly used terms are clarified below. A lifetime incidence, also called a cumulative rate, is calculated using the accepted method of summing the yearly incidences: (Sasieni et al., 2011)

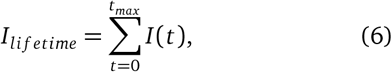

For larger incidence values, the resulting sum produces an exaggerated result. It may become larger than 1 (100%), in which case the use of an approximate clinical statistic called cumulative risk overcomes this issue and is more meaningful. This is much like compound interest, which implicitly assumes an exact exponential incidence progression (Sasieni et al., 2011)

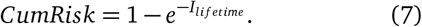

Cumulative risk (Eq. 7) is also an approximation because, in any practical setting, the statistic is complicated by on-going population mortality, multiple diagnoses, and other factors. In addition, cumulative incidence and cumulative risk can be used to find values for any age of interest, not only lifetime. When necessary in this study’s simulations, the exact diagnosis counts were used to calculate the precise cumulative incidence for every age.

### SUPPLEMENTARY FIGURES

**Supplementary Fig. 2.**
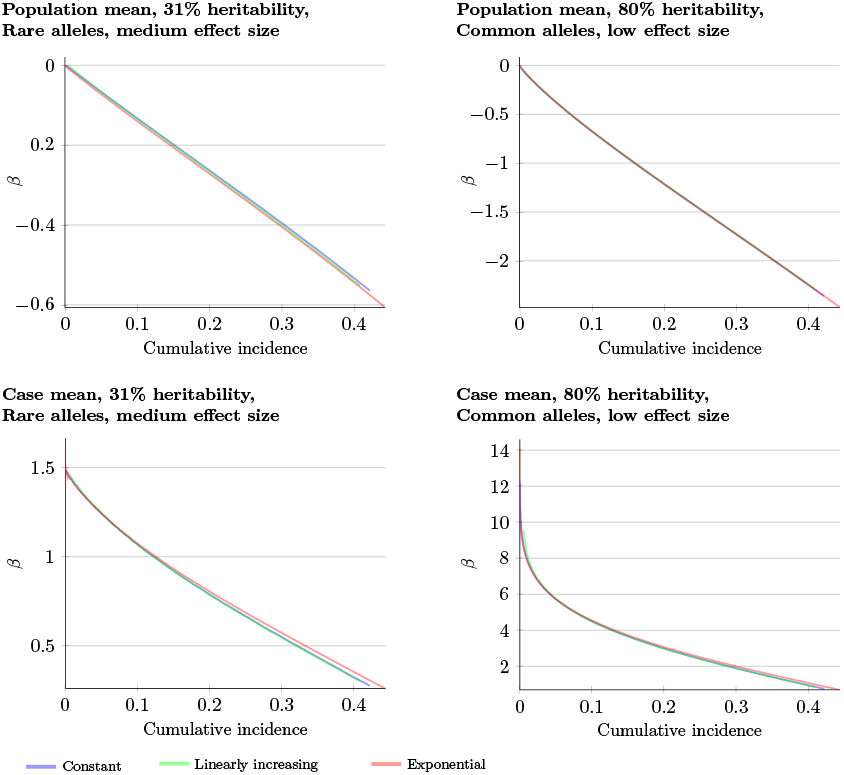
Validation simulations: constant, linear, and exponential incidence curves within the same allele architectures. Using a constant incidence at a level of 0.5% per year, linearly increasing incidence with a slope of 0.01%, and exponentially reaching similar cumulative incidence in a 105-year age interval. Within the same allele architecture, the *β* is identical, subject to the simulation population stochasticity; *β* = *log*(*OddsRatio*).

**Supplementary Fig. 3.**
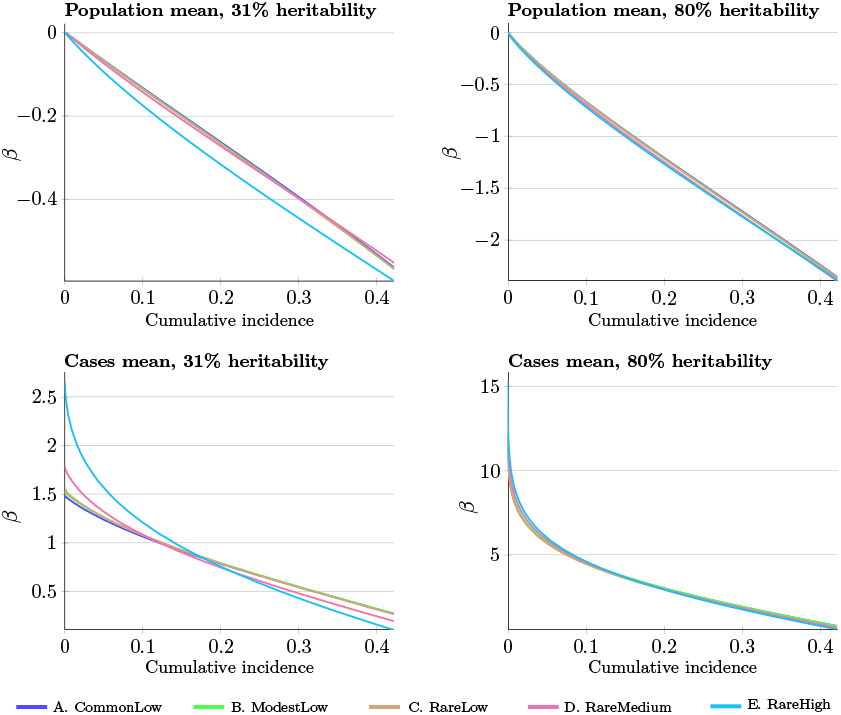
Validation simulations for five allele architectures. The linear and constant incidence patterns give identical results for each allele architecture. The rare medium-effect-size and even rarer high-effect-size scenarios produce a fraction of higher individual betas for the same overall population variance; a relative difference is less prominent at 80% versus 31%. The three identical low-effect-size scenarios produce effectively identical *β* patterns; *β* = *log*(*OddsRatio*).

**Supplementary Fig. 4.**
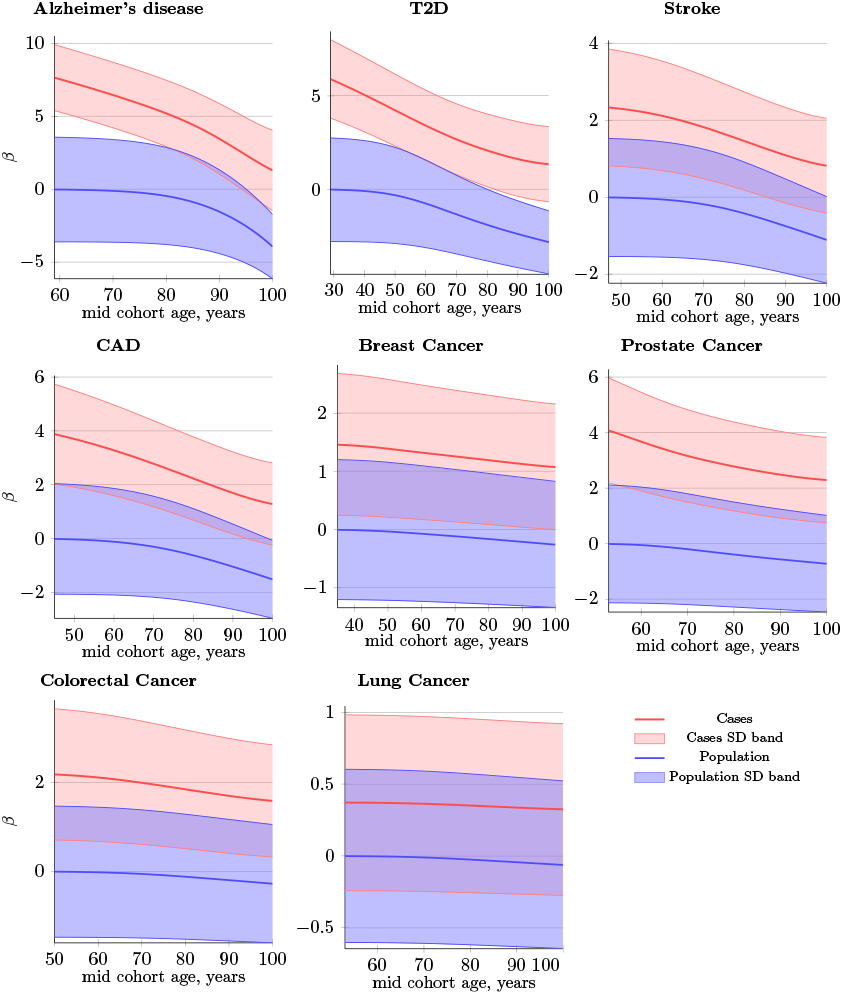
Polygenic score difference between patients and controls in a cohort simulation. Common, low-effect-size alleles (scenario A); *β* = *log*(*OddsRatio*). *SD band* is a band of one standard deviation above and below the cases and the unaffected population of the same age. The cohort change and difference are less prominent than in IVA due to the accumulated diagnoses from younger cases with an averaged control polygenic risk score and mortality.

**Supplementary Fig. 5.**
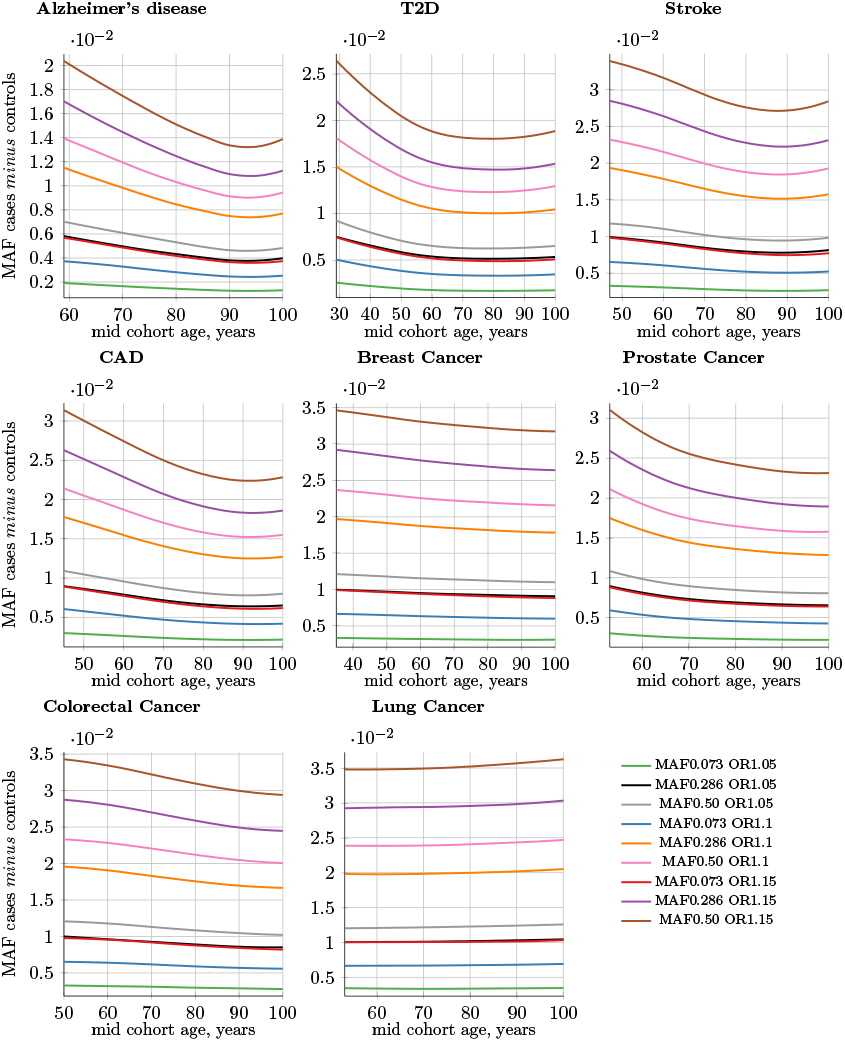
Allele frequency difference between cases and controls; cohort simulation. Common low-effect-size alleles (scenario A). The **MAF cases** *minus* **controls** value is used to determine GWAS statistical power. Rarer and lower-effect-size (OR) alleles are characterized by a lower relative MAF change.

**Supplementary Fig. 6.**
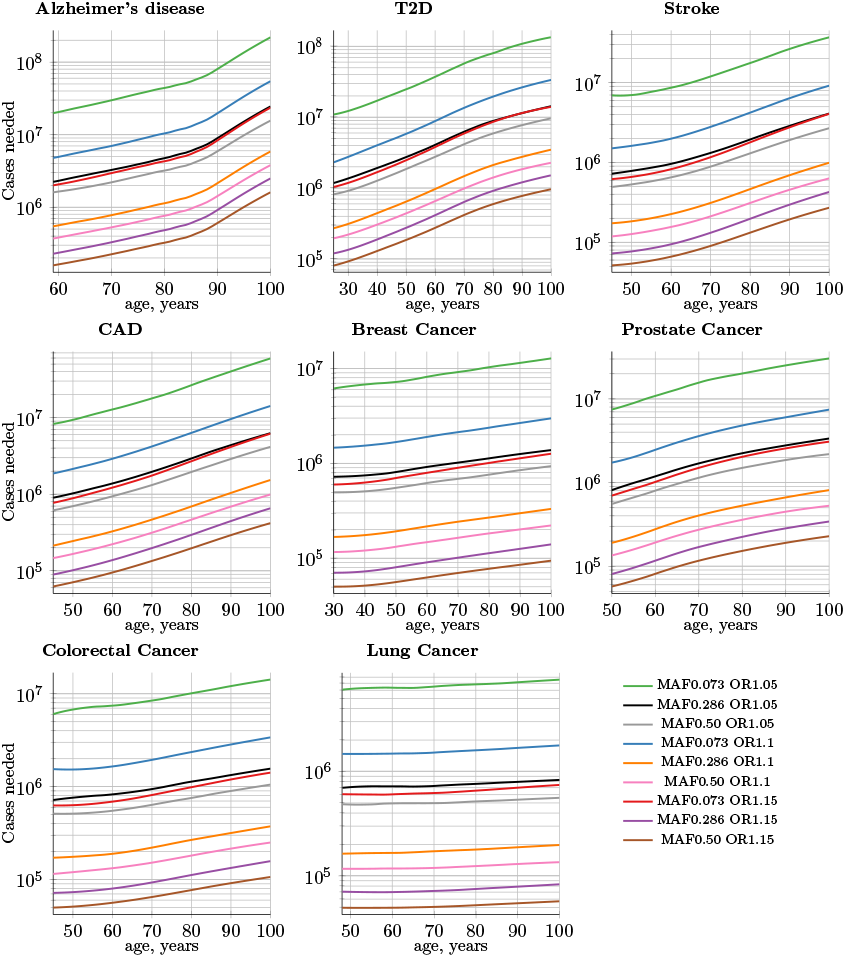
Number of cases needed to achieve 0.8 discovery power; IVA. Common, low-effect-size alleles (scenario A). The diagnosed-individuals-versus-same-age-unaffected-population curve continues to rise steeply in the IVA scenario. A sample of 9 out of 25 SNPs; MAF = minor (risk) allele frequency; OR = risk odds ratio.

**Supplementary Fig. 7.**
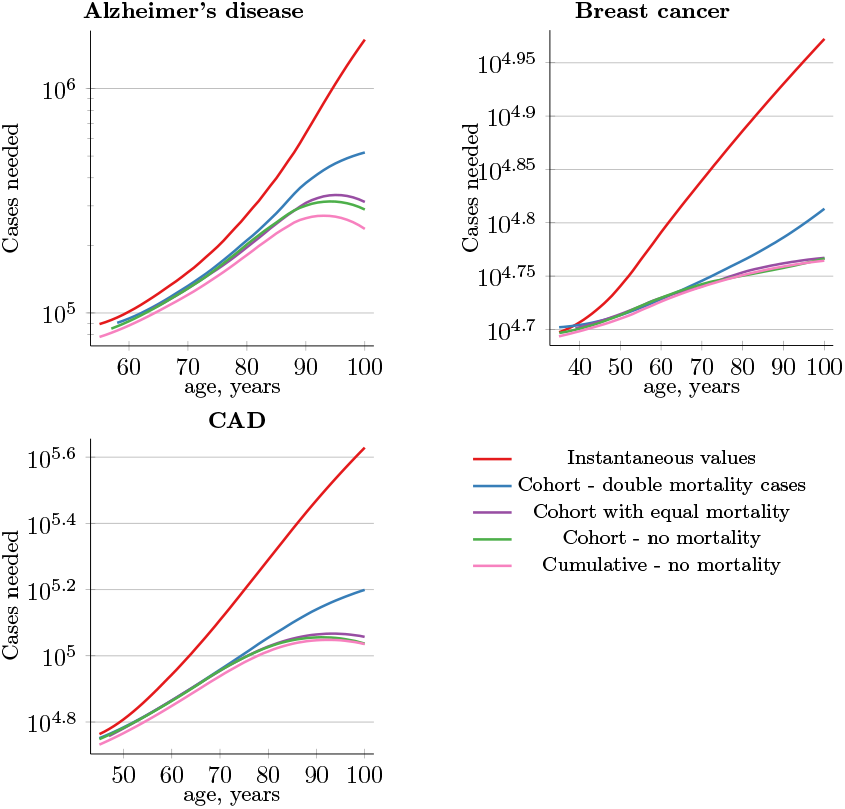
Number of cases needed for 0.8 discovery power for three LODs with representative incidence rate and initial heritability; summary of five LOD validation simulation types. The number of cases needed for 0.8 GWAS discovery power for the clinical cohort study scenario lies between equal mortality for cases and controls and double mortality for cases; it is closer to equal mortality for the LODs we review. The divergence begins after age 85 and is even then relatively modest. “Cohort—double mortality” cases have a mortality twice as large as controls (doubling the value for mortality from the US “Actuarial Life Table”. “Cumulative—no mortality” is the most extreme case of a one-year-span GWAS cohort; with no mortality, it requires the smallest number of cases in GWAS. Note that the logarithmic scale is very different among the three LODs.

**Supplementary Fig. 8.**
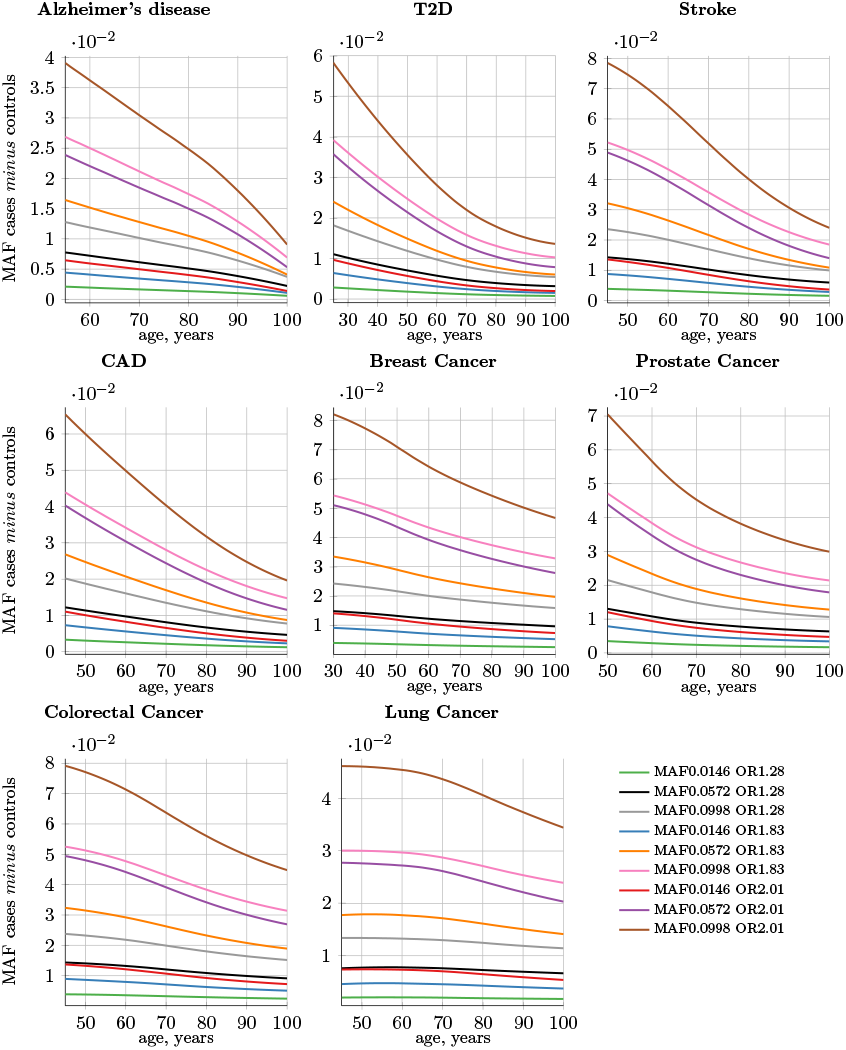
Difference in allele frequency between newly diagnosed instances and the remaining unaffected population; IVA. Rare, medium-effect-size alleles (scenario D). The **MAF cases** *minus***controls** value is used to determine GWAS statistical power. Rarer and lower-effect-size (OR) alleles are characterized by a lower relative MAF change.

**Supplementary Fig. 9.**
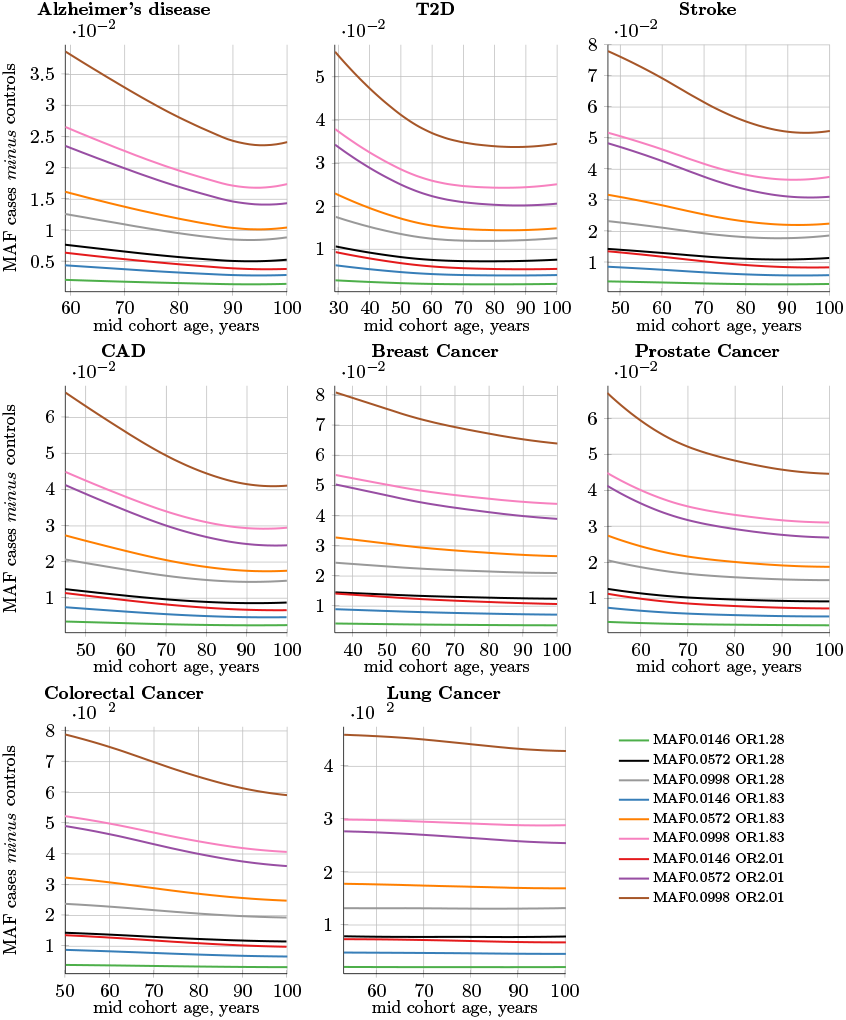
Difference in allele frequency between cases and controls; cohort simulation. Rare, medium-effect-size alleles (scenario D). The **MAF cases** *minus***controlss** value is used to determine GWAS statistical power. Rarer and lower-effect-size (OR) alleles are characterized by a lower relative MAF change.

**Supplementary Fig. 10.**
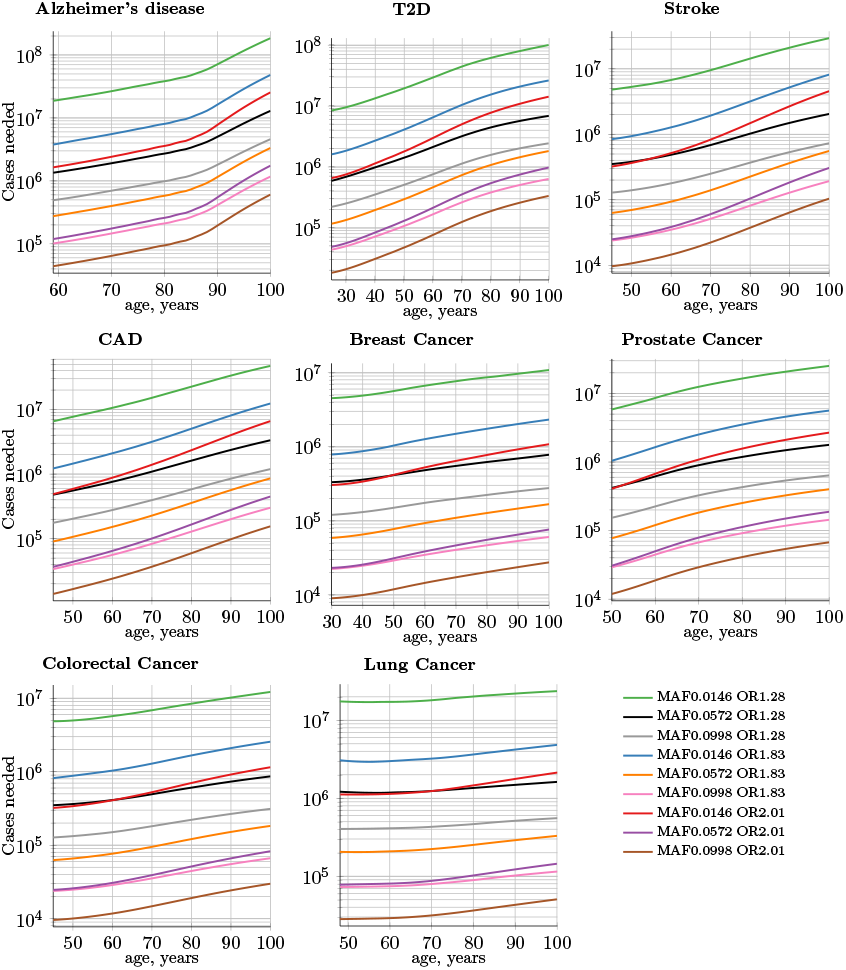
Number of cases needed to achieve 0.8 discovery power; IVA. Rare, medium-effect-size alleles (scenario D). The diagnosed-individuals-versus-same-age-unaffected-population curve continues to rise steeply in the IVA scenario. A sample of 9 out of 25 SNPs; MAF = minor (risk) allele frequency; OR = risk odds ratio.

**Supplementary Fig. 11.**
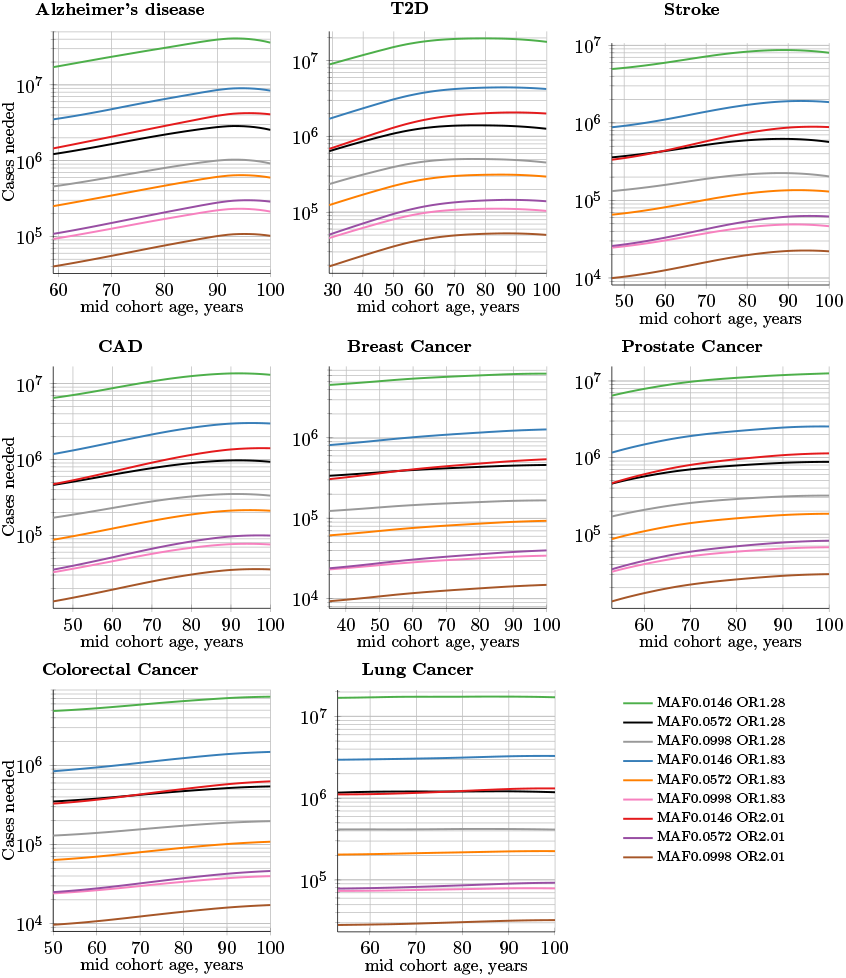
Number of cases needed to achieve 0.8 discovery power; cohort simulation. Rare medium-effect-size alleles (scenario D). The cohort curve due to the accumulative cases diagnosed at younger ages with an averaged control polygenic risk score and mortality begins at the same necessary-cases number as IVA but rises more slowly and levels out at older ages. A sample of 9 out of 25 SNPs; MAF = minor (risk) allele frequency; OR = risk odds ratio.

**Supplementary Fig. 12.**
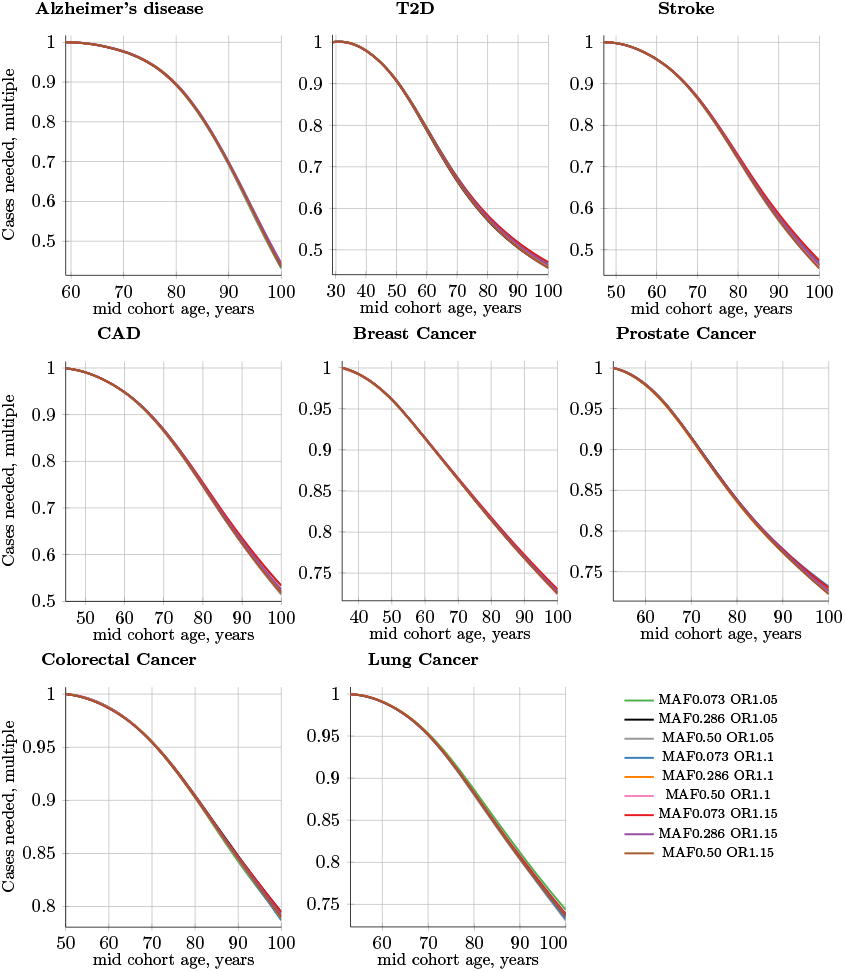
Multiple of the decline in the number of cases needed for 0.8 discovery power in a cohort study using progressively older control cohorts compared to a fixed-age young-cases cohort. Cases’ mid-cohort age is leftmost age (youngest plot point); control mid-cohort ages are incremental ages. The number of cases needed for 0.8 discovery power is smaller when older controls are used, particularly for LODs with the highest heritability and incidence. Common, low-effect-size alleles (scenario A). A sample of 9 out of 25 SNPs; MAF = minor (risk) allele frequency; OR = risk odds ratio.

**Supplementary Fig. 13.**
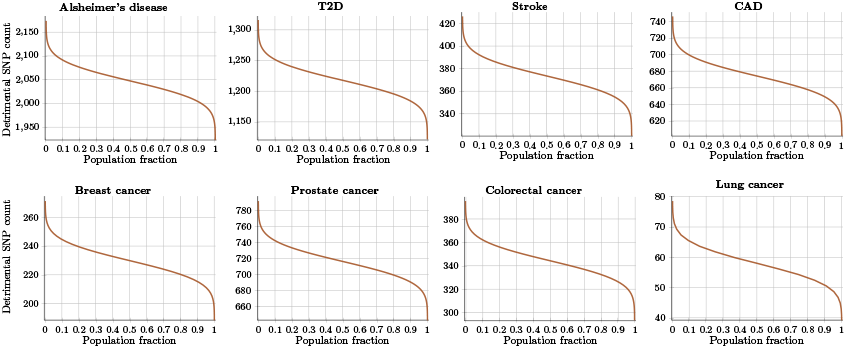
Population distribution of malignant variants for common, low-effect-size genetic architecture. Based on initial heritability, the individuals in a population carry a relatively high number of malignant, low-effect alleles, resulting in the combined LOD PRS.

**Supplementary Fig. 14.**
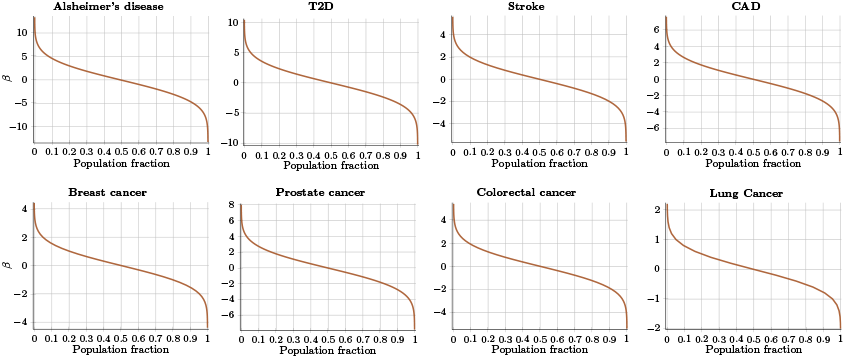
Population distribution of PRSs for common, low-effect-size genetic architecture. *β* = *log*(*OddsRatio*) normalized to population mean.

**Supplementary Fig. 15.**
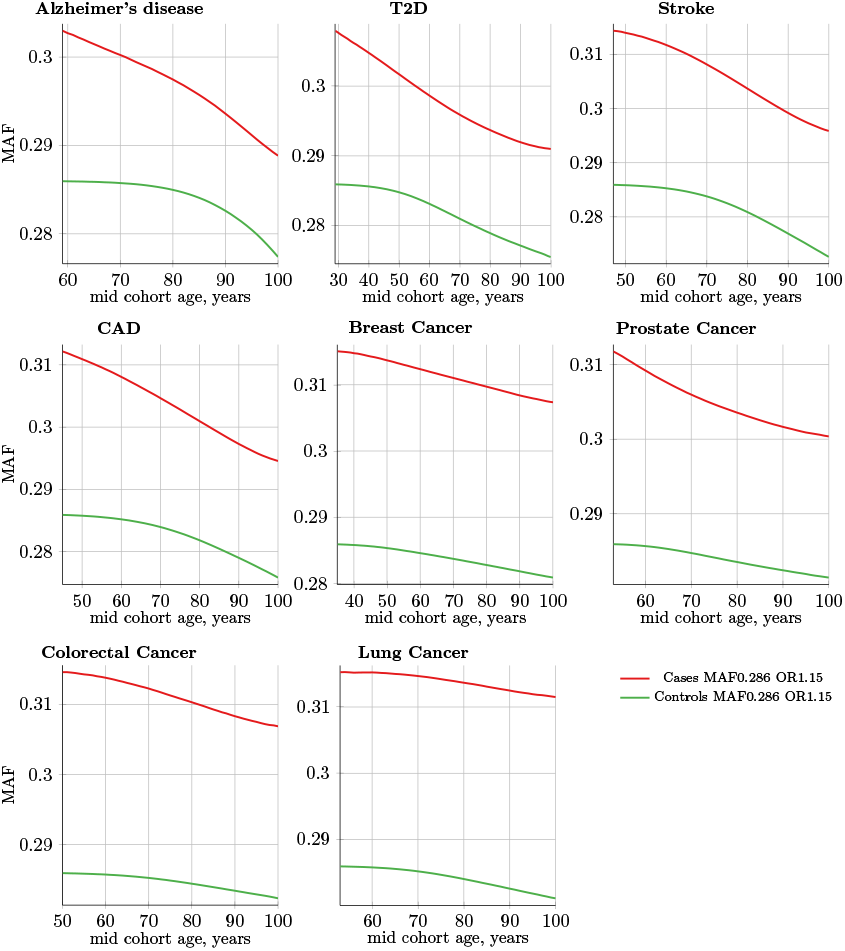
Absolute magnitude change in MAF (minor allele frequency) with age for cases and controls; cohort simulation. Common, low-effect-size alleles (scenario A), all plots show MAF = 0.286 and OR = 1.15 allele. Change in the absolute magnitude of each allele frequency value is relatively small with age progression. GWAS discovery power is a function of the difference in allele frequency between cases and controls. Rarer and lower-effect-size (OR) alleles are characterized by a lower change in absolute and relative MAF with cohort age progression.

**Supplementary Fig. 16.**
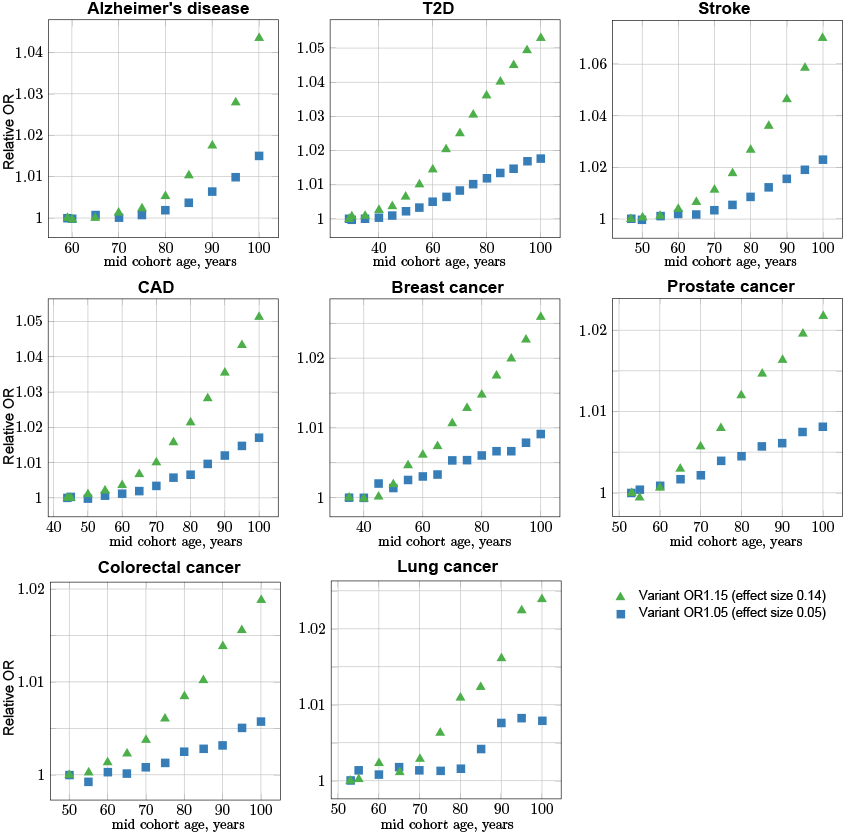
GWAS association simulations: OR bias progression with control cohort age increasing against the constant youngest possible case cohort. Common, low-effect-size alleles (scenario A), showing two SNPs—the with the largest and the smallest effect—for each LOD. The OR increase (bias) with mid-cohort age progression implies a power of ΔAge from age matched youngest cohort. The confidence intervals are not displayed on this plot for illustration purpose; they are displayed in Supplementary Fig. 17, showing the same data in effect size units.

**Supplementary Fig. 17.**
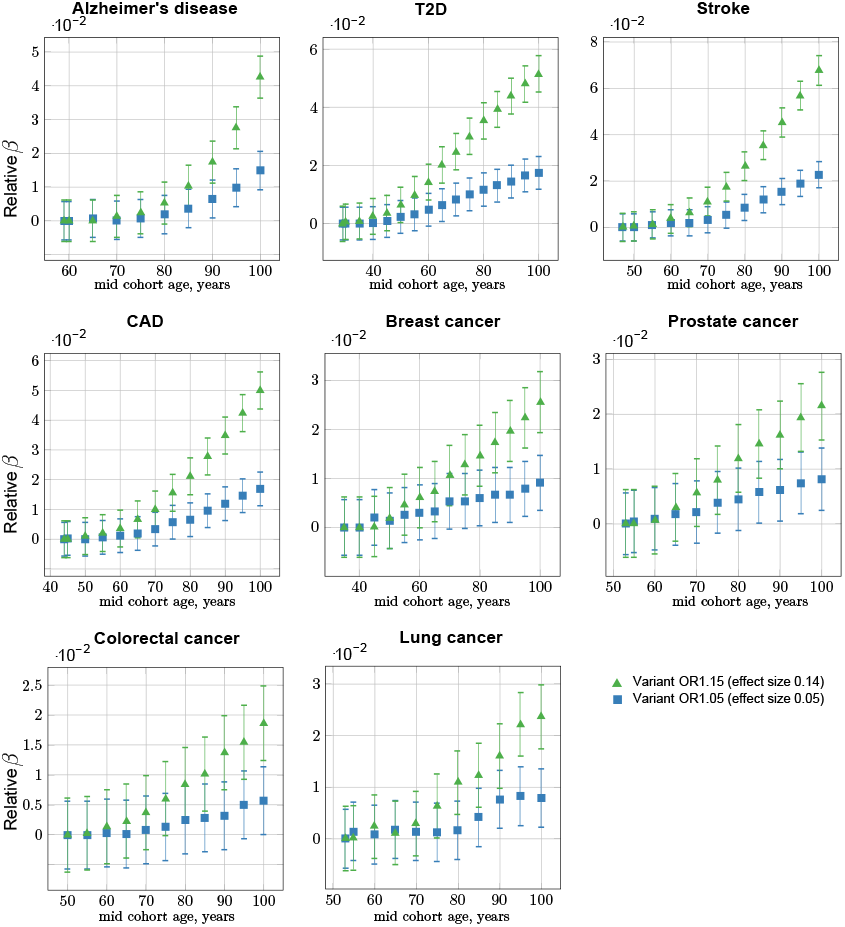
GWAS association simulations: relative effect size bias progression with control cohort age increasing against the constant youngest possible case cohort. Common, low-effect-size alleles (scenario A), showing two SNPs—the with the largest and the smallest effect—for each LOD. The confidence interval bars correspond to two-sigma (95%) confidence from the GWAS logistic regression association. The OR increase with mid-cohort age progression implies a power law relative to Δ*age*. This plot implies the LOD SNP age bias and corresponding adjustment value proportionate to the SNP effect size; this observation is further investigated in Supplementary Fig. 18 and Supplementary Fig. 19.

**Supplementary Fig. 18.**
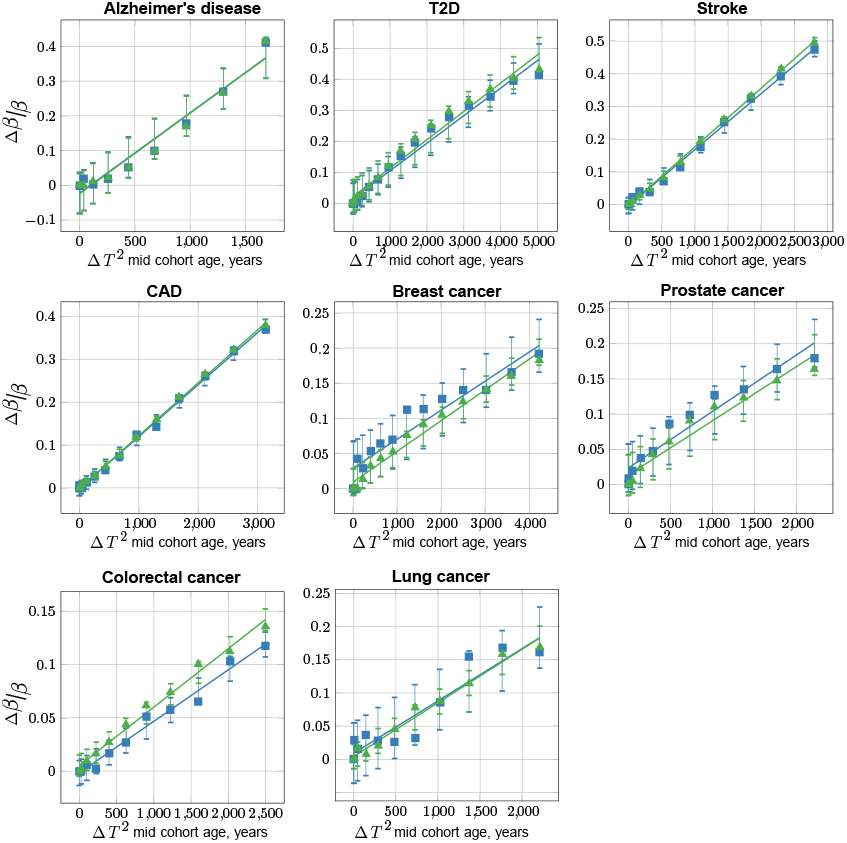
GWAS association simulations: characterizing the age bias adjustment maintaining “true" OR with control cohort age progression (quadratic: Δ*T*^2^). Common, low-effect-size alleles (scenario A), showing two SNPs—the with the largest and the smallest effect—for each LOD. The confidence interval bars correspond to two-sigma (95%) based on standard error of linear regression fitting (drawn relative to the regression line). This plot depicts the adjustment proportionate to square of Δ*t* = *t* − *T*_Y_-relative age from the youngest cohort mid-cohort age for the normalized bias of the effect size *β* calculated Δ*β*/*β*, as described in the main article. See also Supplementary Table 8.

**Supplementary Fig. 19.**
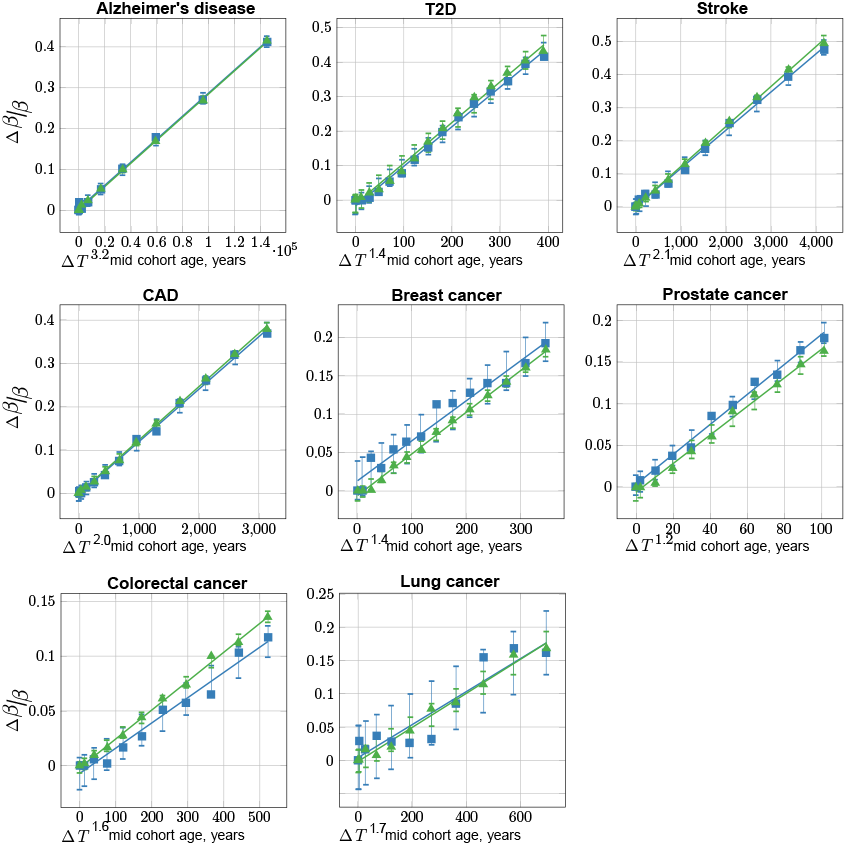
GWAS association simulations: characterizing the age bias adjustment maintaining “true” OR with control cohort age progression (best fit power: Δ*T*^*P*^). Common, low-effect-size alleles (scenario A), showing two SNPs—the with the largest and the smallest effect—for each LOD. The confidence interval bars correspond to two-sigma (95%) based on standard error of linear regression fitting (drawn relative to the regression line). In this plot, rather than using the square of Δ*age*, the best fit power is iteratively discovered, achieving better residual standard error and P-value of the R lm() regression, compared to Supplementary Fig. 18.

**Supplementary Table 8.** Summary of GWAS association simulations and age covariate correction parameters for youngest cases—older controls cohorts.

| Disease | Highly prevalent LODs |  |  |  | Cancers |  |  |  |
| --- | --- | --- | --- | --- | --- | --- | --- | --- |
|  | AD | T2D | Stroke | CAD | Breast | Prostate | Colorectal | Lung |
| Youngest cases mid-cohort age | 59 | 29 | 47 | 44 | 35 | 53 | 50 | 53 |
| **GWAS Association SSE for $\beta = 0.14$ (OR1.15):** |  |  |  |  |  |  |  |  |
| 100Y controls $\beta$ SSE raw | 0.00312 | 0.00311 | 0.00312 | 0.00311 | 0.00310 | 0.00310 | 0.00310 | 0.00310 |
| 100Y controls $\beta$ SSE adjusted | 0.00342 | 0.00345 | 0.00359 | 0.00336 | 0.00315 | 0.00314 | 0.0312 | 0.00314 |
| **GWAS Association SSE for $\beta = 0.05$ (OR1.05):** |  |  |  |  |  |  |  |  |
| 100Y controls $\beta$ SSE raw | 0.00283 | 0.00283 | 0.00283 | 0.00283 | 0.00283 | 0.00283 | 0.00283 | 0.00283 |
| 100Y controls $\beta$ SSE adjusted | 0.00310 | 0.00311 | 0.00321 | 0.00304 | 0.00288 | 0.00288 | 0.00285 | 0.00287 |
| **Age bias adjustment - quadratic ($\Delta age^2$):** |  |  |  |  |  |  |  |  |
| Slope coefficient | $2.3 \cdot 10^{-4}$ | $9.1 \cdot 10^{-5}$ | $1.8 \cdot 10^{-4}$ | $1.2 \cdot 10^{-4}$ | $4.4 \cdot 10^{-5}$ | $7.7 \cdot 10^{-5}$ | $5.5 \cdot 10^{-5}$ | $8.1 \cdot 10^{-5}$ |
| Residual standard error | 0.029 | 0.026 | 0.0058 | 0.0039 | 0.0093 | 0.014 | 0.0050 | 0.0092 |
| P-value | $5.5 \cdot 10^{-7}$ | $1.9 \cdot 10^{-12}$ | $2.9 \cdot 10^{-16}$ | $2.7 \cdot 10^{-18}$ | $1.8 \cdot 10^{-11}$ | $4.1 \cdot 10^{-7}$ | $2.7 \cdot 10^{-10}$ | $5.3 \cdot 10^{-9}$ |
| **Age bias adjustment - best fit power ($\Delta age^P$):** |  |  |  |  |  |  |  |  |
| Power | 3.2 | 1.4 | 2.1 | 2.0 | 1.4 | 1.2 | 1.6 | 1.7 |
| Slope coefficient | $2.8 \cdot 10^{-6}$ | $1.2 \cdot 10^{-3}$ | $1.2 \cdot 10^{-4}$ | $1.2 \cdot 10^{-4}$ | $5.4 \cdot 10^{-4}$ | $1.7 \cdot 10^{-3}$ | $2.6 \cdot 10^{-4}$ | $2.6 \cdot 10^{-3}$ |
| Residual standard error | 0.0030 | 0.013 | 0.0057 | 0.0039 | 0.0036 | 0.0053 | 0.0025 | 0.0084 |
| P-value | $6.9 \cdot 10^{-15}$ | $1.3 \cdot 10^{-16}$ | $2.5 \cdot 10^{-16}$ | $2.7 \cdot 10^{-18}$ | $2.0 \cdot 10^{-16}$ | $5.3 \cdot 10^{-11}$ | $5.8 \cdot 10^{-13}$ | $2.4 \cdot 10^{-9}$ |

The two *GWAS association...* sections summarize the standard error of the logistic regression association for cohort studies with the largest age difference between youngest case group (as specified for each LOD in this table) and a control group with the mid-cohort age of 100 years. The “raw” value corresponds to the analysis without the age bias adjustment, and “adjusted” - after the age bias adjustment. The two *Age bias adjustment...* sections show parameters of the regression described in the article’s Methods section when using quadratic and best fit power for the age difference between the youngest cases mid-cohort age and incrementally increasing mid-cohort age of older controls for the SNP with “true” OR=1.15. The best fit power results in a more accurate regression, yet the quadratic rule would be sufficiently accurate for the practical GWASs data.

